# Lysosomal Degradation Pathways Target Mutant Calreticulin and the Thrombopoietin Receptor in Myeloproliferative Neoplasms

**DOI:** 10.1101/2023.07.12.548605

**Authors:** Amanpreet Kaur, Arunkumar Venkatesan, Malathi Kandarpa, Moshe Talpaz, Malini Raghavan

**Affiliations:** Department of Microbiology and Immunology, University of Michigan Medical School, Ann Arbor, MI, USA, 48109; Department of Ophthalmology & Visual Sciences, Upstate Medical University, Syracuse, NY, USA, 13202; Department of Internal Medicine, Division of Hematology/Oncology, University of Michigan Rogel Cancer Center, Ann Arbor, MI, USA, 48109

**Keywords:** Myeloproliferative Neoplasms, calreticulin, MPL, thrombopoietin receptor, lysosomal degradation, rapamycin, everolimus, mTOR inhibitors

## Abstract

Somatic mutants of calreticulin (CRT) drive myeloproliferative neoplasms (MPNs) via binding to the thrombopoietin receptor (MPL) and aberrant activation of the JAK/STAT pathway. Compared with healthy donors, platelets from MPN patients with CRT mutations display low cell surface MPL. Co-expression of MPL with an MPN-linked CRT mutant (CRT_Del52_) reduces cell surface MPL expression, indicating the involvement of induced protein degradation, a better understanding of which could lead to new therapies. We show that lysosomal degradation is relevant to the turnover of both CRT_Del52_ and MPL. Drug-mediated activation of lysosomal degradation reduces CRT_Del52_ and MPL expression, with parallel inhibition of CRT_Del52_-induced cell proliferation and stem cell colony formation. Thus, reduced surface MPL, a marker of platelets from MPN patients with CRT mutations, results from mutant CRT-induced lysosomal degradation of MPL. Drug-induced activation of lysosomal degradation compromises the pathogenic effects of CRT_Del52_, which can be further exploited for therapeutic interventions.

## Introduction

Essential thrombocythemia (ET), polycythemia vera (PV) and myelofibrosis (MF) are myeloproliferative neoplasms (MPNs) that include hyperproliferation of myeloid lineage blood cells ^1, 2^. The molecular mechanism underlying the development of these cancers involves a set of driver mutations that result in constitutive hyperactivation of signaling pathways that regulate hematopoiesis ^1, 2^. Somatic mutations of the calreticulin (CRT) gene are found in 20-35 % of ET and MF patients ^3, 4^. Most of these mutations are +1 bp frame shift mutations in the exon 9 that encodes the carboxy domain of CRT. The type 1 mutation, which involves a 52 bp deletion (Del52), and the type 2 mutation, which involve a 5 bp insertion (Ins5), account for ∼45 to 53% and ∼32 to 41% of CRT mutations in MPNs, respectively ^3, 4^. MPN-linked mutant CRT proteins are characterized by enrichment of basic amino acids in the carboxy domain in contrast to acidic amino acid residues present in the wild-type CRT protein ^3, 4^. The MPN CRT mutants also lack the KDEL motif at the carboxy-terminal end which is responsible for endoplasmic reticulum (ER) localization of the wild-type protein. Loss of the KDEL sequence affects ER retention ^3–5^ and induces secretion of MPN-linked CRT mutants ^6–9^.

The pathogenic effects of mutant CRT proteins in MPNs are attributed in part to their ability to induce ligand-independent constitutive activation of JAK/STAT signaling via the cell surface receptor, MPL (also called the thrombopoietin receptor (TPOR)), that is also a glycoprotein substrate of CRT ^10–14^. While the interaction between MPL and wild-type CRT is transient, mutant CRT proteins form stable complexes with MPL via both their glycan-binding site ^5, 14–16^ and the novel C-terminal domain ^15, 17, 18^. The mutant CRT and MPL complexes co-traffic to the cell surface ^5^. MPL is expressed on the surface of hematopoietic cells and regulates the differentiation of the megakaryocyte lineage and production of platelets as well as self-renewal of HSCs ^19–21^. Expression of MPN-linked mutant CRT proteins and the resultant activation of MPL signaling, promotes proliferation of megakaryocytes and excessive production of platelets ^10, 11, 13, 22–24^.

The physiological activation of MPL signaling by TPO is tightly regulated via coordination between the synthesis and release of TPO and its removal from blood circulation ^25^. TPO is synthesized within the liver and to a small extent within the kidneys and released into the blood circulation ^26^. Binding of TPO to MPL and the resultant activation of receptor signaling is followed by internalization and degradation of the receptor-ligand complex which, in turn, removes excess TPO from circulation ^27–30^. However, little is known about the downstream effects of MPL signaling activated by the binding of MPN-linked mutant CRT proteins. In this study, we measured the relative levels of MPL protein on platelets purified from MPN patients and healthy donors and found a significant reduction in MPL levels in MPN patient platelets. We used model cell lines co-expressing human MPL and mutant CRT protein to explore pathways involved in the regulation of surface MPL and mutant CRT levels. Our studies provide insights into the pathways mediating mutant CRT and MPL degradation that can be exploited for the treatment of MPNs.

## Results

### Platelets from MPN patients are characterized by surface expression of mutant calreticulin

The cell surface localization of MPN-linked CRT mutants has been seen earlier on lymphocytes and monocytes ^8^ as well as on CD34^+^ hematopoietic stem cells and CD34^+^ cell-derived megakaryocytes ^16^ from mutant CRT expressing MPN patients. We have previously detected the expression of mutant CRT proteins in the lysates of platelets isolated from MPN patients ^17^. In this study, we examined the localization of CRT proteins on the surface of platelets isolated from blood samples of MPN patients (details of patient samples, disease and mutation are in Supplemental Table 2) or healthy donors that were concurrently processed. The mutant CRT proteins were detected using a mutant specific antibody (anti-CRT(C_mut_)) directed against 22 amino acids unique to the carboxy domain sequence of MPN-linked mutant CRT proteins ^17^. We also used a commercial antibody that binds to the non-mutated amino terminal globular domain (anti-CRT(N)) of CRT protein present in both the wild type and mutant CRT proteins. Using flow cytometry, platelets were gated on single cells followed by gating for CD41 (GPIIb) present on the surface of platelets in a heterodimeric complex with CD61 (GPIIIa) ^31^. CD62p (P selectin), can be used to identify activated platelets ^32, 33^.

We first examined CD41 and CRT staining patterns in healthy donor platelets by flow cytometry following staining of cells without permeabilization (surface) or after membrane permeabilization with 0.2% saponin (total). Setting detectable protein levels in the permeabilized cells at 100%, a considerable fraction of the total CD41 is detected on the cell surface (**Figure 1B**), which agrees with the well-studied surface localization of platelet CD41 protein. Calreticulin, on the other hand, is predominantly ER localized, but known to translocate to cell surface under certain conditions ^34^. Our results show low background level staining of CRT protein with the anti-CRT(N) antibody in non-permeabilized cells (surface) compared with permeabilized cells (total), confirming the predominant intracellular localization of CRT in platelets (**Figure 1C**). Compared to healthy donor platelets, we observed significantly higher mean fluorescence intensity (MFI) values of surface staining with both anti-CRT(C_mut_) (**Figure 1D**) and anti-CRT(N) (**Figure 1E**) antibodies on CD41^+^ platelets from MPN patients. A similar trend was also noted on CD62p^+^ (activated) platelets from MPN patients compared to healthy control platelets (**Figures 1F and G**). These results demonstrate that MPN-linked mutant CRT proteins, that lack ER retention signals, are detectable on the surface of platelets isolated from MPN patients.

**Figure 1:**
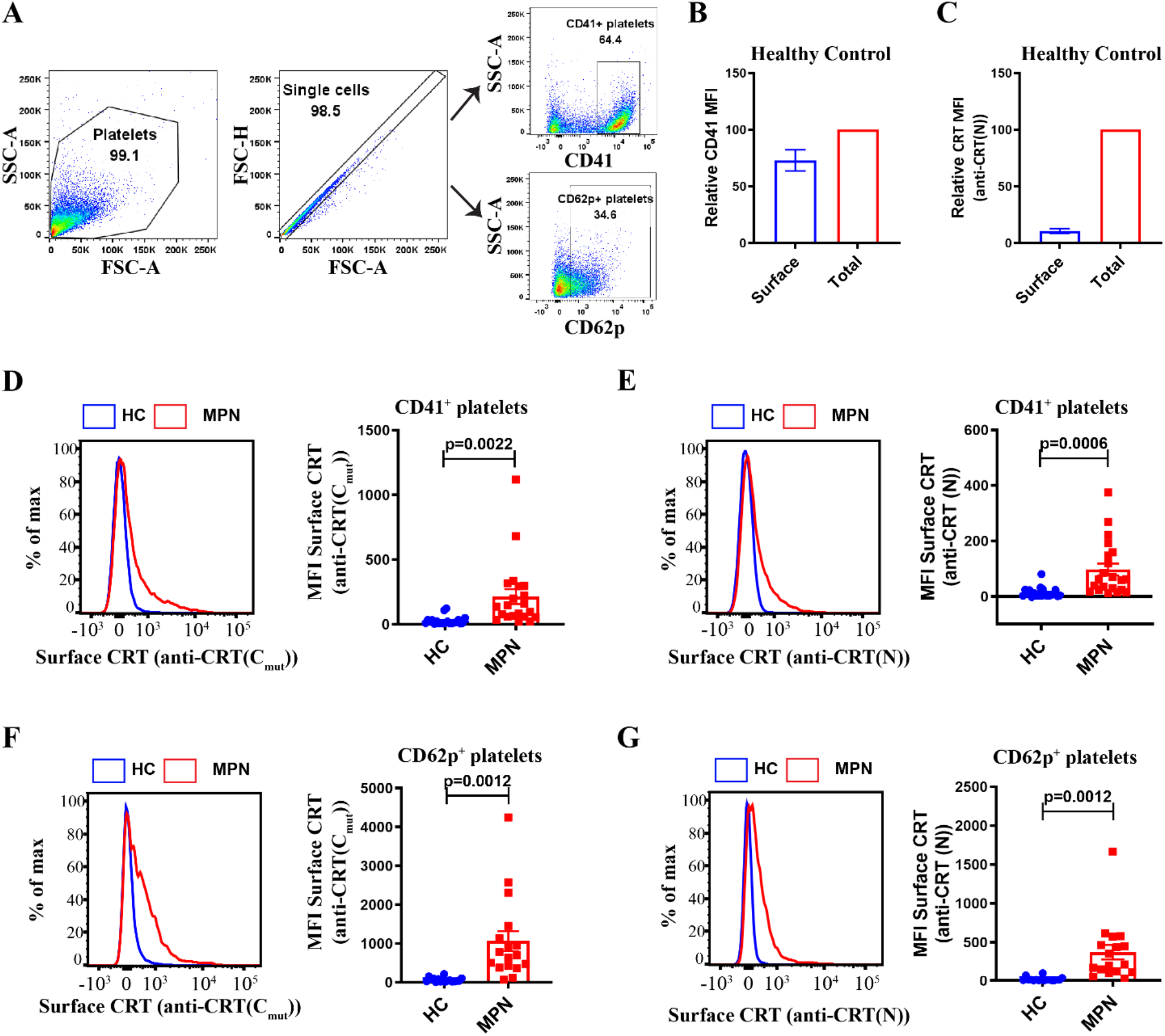
Platelets from MPN patients are characterized by cell surface expression of mutant calreticulin: Platelets were isolated from blood samples of mutant CRT expressing MPN patients and healthy controls. **(A)** Gating strategy used for the analysis of CD41^+^ and CD62p^+^ platelets by flow cytometry. **(B-C)** Bar graphs indicate flow cytometry-based detection of staining for CD41 **(B)** and CRT (detected using an N domain specific antibody anti-CRT(N)) **(C)** on healthy donor platelets either after fixation alone (surface) or after fixation and permeabilization (total). **(D-G)** Flow cytometry-based detection of surface CRT on CD41^+^ **(D&E)** and CD62p^+^ **(F&G)** platelets. Representative histograms and bar graphs show mean fluorescence intensity (MFI) of surface CRT detected on platelets by either mutant CRT specific antibody (anti-CRT (C_mut_)) or anti-CRT (N) antibody. MPN patient and healthy donor (HC) platelet samples that were processed together were analyzed as pairs for these measurements (n=21 for measurements on CD41^+^ platelets and n=17 for CD62p^+^ platelets). The error bars show the standard error mean (SEM). Statistical significance indicated by p values was determined using GraphPad Prism and paired t-test analyses.

### Low surface MPL levels in MPN patient platelets

Feedback regulation of MPL signaling via its degradation is important for keeping a check on the proliferation of hematopoietic cells ^27, 29, 30^. However, whether this feedback control regulates the ligand-independent constitutive activation of MPL signaling by mutant CRT proteins is unknown. We measured the levels of MPL protein on platelets isolated from MPN patients expressing mutant CRT proteins. The surface MPL levels were measured on CD41^+^ platelets using a commercial anti-MPL polyclonal antibody (anti-CD110 clone 1.6.1) ^35^. Platelets isolated from the majority of MPN patients showed lower surface MPL levels than those on the healthy donor platelets, suggesting a specific downmodulation of surface MPL in MPN patient platelets that overrides normal MPL expression variations that may exist among individuals (**Figures 2A and 2B**). To determine if the reduction in surface MPL is also reflected in the levels of total MPL protein in MPN patient platelets, we stained platelets for total MPL protein following permeabilization of platelets. Again, CD41^+^ platelets isolated from most of the patients exhibited low total MPL levels compared to healthy donor platelets (**Figures 2C and 2D**). Platelet lysates from MPN patients also exhibit a trend towards lower MPL protein levels detected by immunoblotting compared to the levels in the lysates of healthy donor platelets (**Figure 2E**). Thus, platelets from MPN patients expressing CRT mutants exhibit low surface and total MPL protein levels, which is a potential biomarker for mutant CRT^+^ MPN.

**Figure 2:**
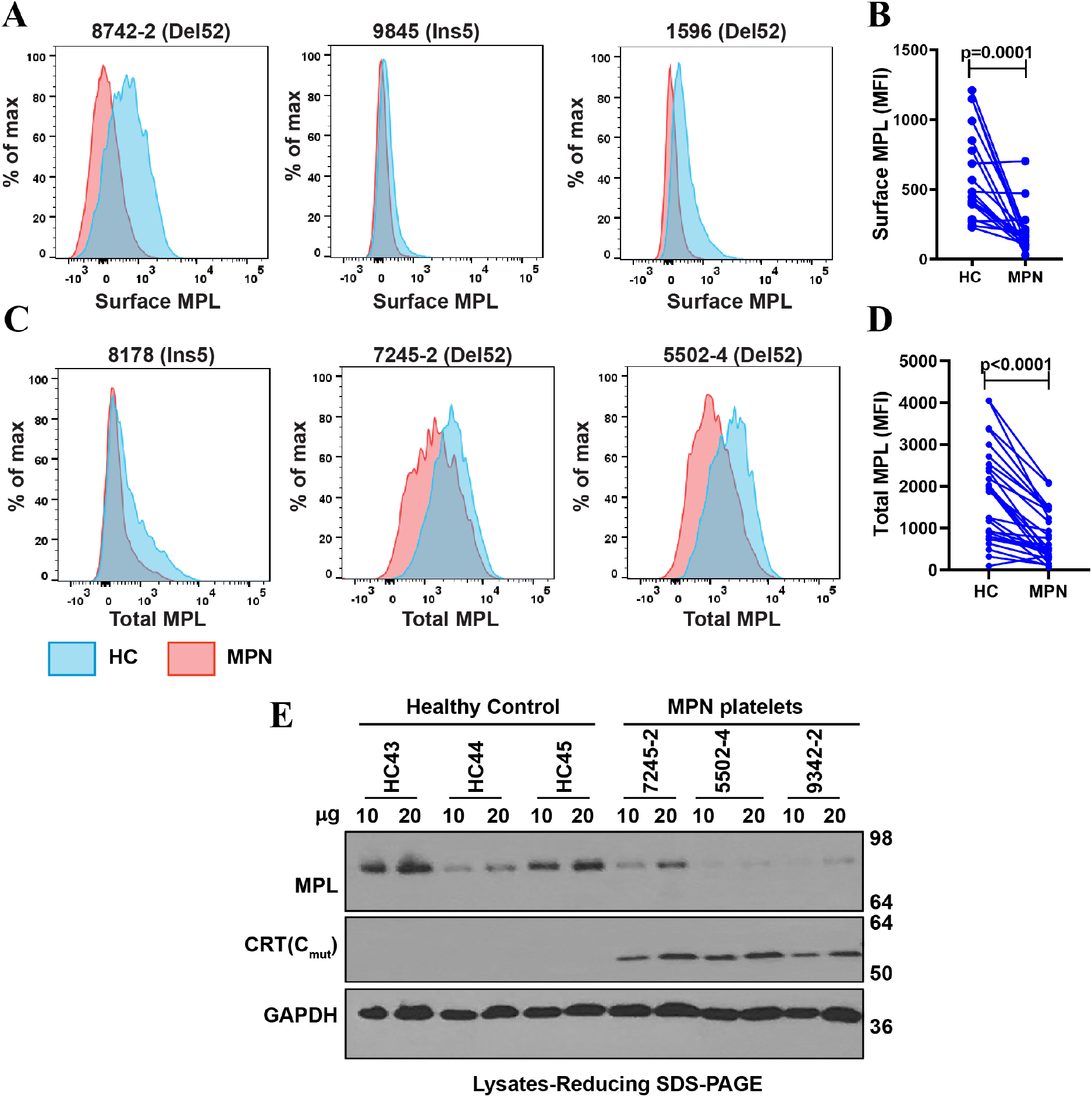
Downmodulation of MPL in MPN patient platelets: (A and C) Representative histograms show surface (A) or total (C) MPL staining with anti-MPL antibody on MPN patient (MPN) platelets (red histograms) or same day healthy control (HC) platelets (blue histograms) in non-permeabilized (A) or permeabilized (C) platelets. **(B and D)** The mean fluorescence intensity values of surface (B; n=19) and total (D; n=28) MPL staining on platelets are shown here for pairs of MPN patient and healthy control (HC) samples that were processed simultaneously. Each dot in **(B)** and **(D)** represents the MFI value for an individual MPN or HC platelet sample, while the lines connect pairs of HC and MPN patient samples that were processed simultaneously. Statistical significance indicated by p values was determined using GraphPad Prism and a paired t-test analysis. **(E)** Representative blots showing MPL and mutant CRT protein levels in the lysates of healthy control or MPN patient platelets subjected to SDS-PAGE under reducing conditions followed by immunoblotting. Anti-MPL and anti-CRT(C_mut_) antibodies were used for detection of MPL and mutant CRT proteins, respectively. GAPDH blot shows relative loading of lysates.

### Reduction of total and surface human MPL levels when co-expressed with CRT_Del52_ in Ba/F3 cells

To further investigate whether surface MPL downmodulation is induced by co-expression of MPN-linked CRT mutants, we expressed human MPL protein in the murine pro-B cell line, Ba/F3, together with CRT_Del52_, a frequently occurring type I MPN CRT mutant ^3, 4^. Ba/F3 cells require cytokine (IL3) for growth and have been used in various studies of MPN-linked CRT mutants. Ba/F3 cells were transduced with retroviruses for stable expression of human MPL protein either alone (empty vector control (EV)) or together with either CRT_WT_ or CRT_Del52._ We observed a significant downmodulation of surface and total MPL protein levels in Ba/F3-MPL CRT_Del52_ cells compared to the CRT_WT_ expressing or EV cells (**Figures 3A-C**).

**Figure 3:**
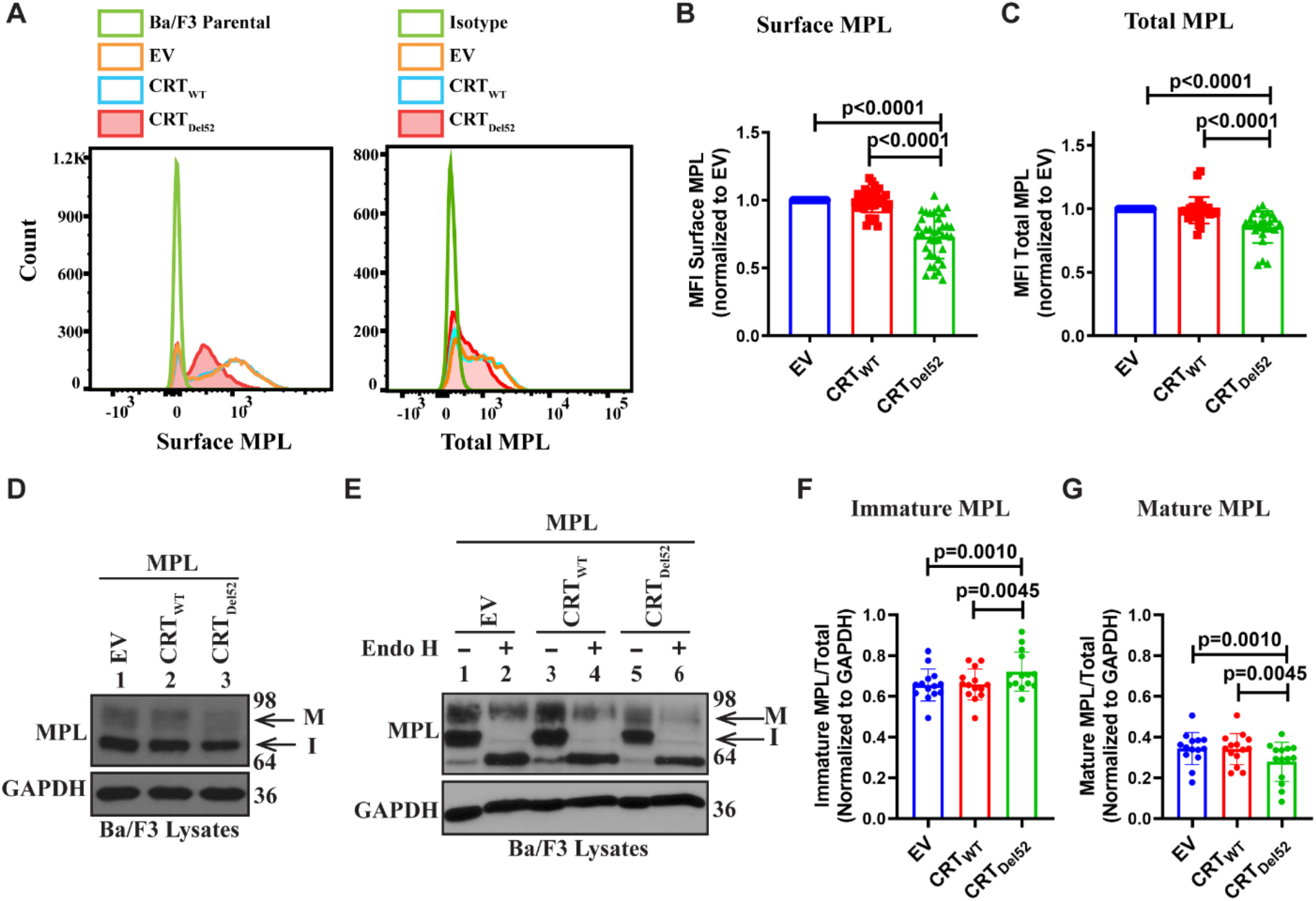
Reduction of total and surface human MPL levels when co-expressed with CRT_Del52_ in Ba/F3 cells: IL-3 dependent murine Ba/F3 cells were transduced for co-expression of human MPL and either WT human CRT (CRT_WT_) or mutant human CRT (CRT_Del52_) protein. Ba/F3 cells expressing only human MPL protein (Empty Vector, EV) were included as controls. **(A)** Representative histograms show surface and total MPL staining detected by flow cytometry in Ba/F3-MPL cells expressing either MPL alone or MPL along with CRT_WT_ or CRT_Del52_ as indicated. **(B)** Bar graphs show quantifications of mean fluorescence intensities of surface MPL **(B)** and total MPL **(C)** measured by flow cytometry in CRT_WT_ or CRT_Del52_ expressing cells normalized to the MFI values of EV cells within the same experiments. The data points represent measurements taken in 38 (B) and 25 (C) independent experiments using cells from ∼8 different retroviral infections. P-values show statistical significance determined by paired t-test analyses using GraphPad Prism. **(D and E)** Representative immunoblots (D, n=14 and E, n=3) showing the levels of total MPL protein in Ba/F3 cells co-expressing either CRT_Del52_ or CRT_WT_ compared to empty vector (EV) control cells (D) and without or with Endo H digestion (E). GAPDH is shown for loading controls. Top and bottom bands in the MPL blots represent the mature (M) and immature (I) forms of MPL proteins, respectively. **(F and G)** Bar graphs show fraction of immature (F) and mature (G) human MPL protein in EV, CRT_WT_ and CRT_Del52_ expressing cells quantified from intensities of the mature and immature MPL bands from the immunoblots using image J software normalized to the intensities of GAPDH bands. Paired t-tests were used in GraphPad Prism to determine statistical significance, indicated by p-values.

We further checked the MPL protein levels in lysates of Ba/F3 cells expressing MPL alone (EV) or those co-expressing human MPL with either CRT_WT_ or CRT_Del52_. Two distinct forms of MPL protein are detected (with anti-MPL antibody- 06944 from EMD Millipore) in the lysates upon SDS-PAGE, as previously described ^36^. Endoglycosidase H (Endo H) is an enzyme that cleaves immature N-glycans rich in mannose residues acquired by glycoproteins in the ER (Endo H-sensitive) but not complex glycans (Endo H-resistant) that have been processed by enzymes within the Golgi complex during glycoprotein trafficking to cell surface. The slower migrating MPL band (just below the 98 kDa marker in **Figures 3D and 3E**) represents the mature MPL protein (M), based on its resistance to Endo H digestion. The faster migrating band (halfway between the 98kDa and 64 kDa markers in **Figures 3D and 3E**) represents the immature MPL (I) band, as Endo H digestion causes its shift to a lower molecular weight (**Figure 3E**). We observe an increase in the fraction of immature MPL in CRT_Del52_ expressing cells (**Figure 3F**) with a concomitant decrease in the mature protein fraction (**Figure 3G**) when compared to EV or CRT_WT_ expressing cells. This result is consistent with the observed downmodulation of cell surface MPL in platelets from MPN patients and Ba/F3 cells (Figures 2 and 3A, B) as cell surface MPL is expected to be largely Endo H resistant.

### Lysosomal degradation of surface/mature MPL is enhanced in the presence of CRT_Del52_

We next assessed whether altered protein degradation might explain low surface/mature MPL levels in CRT_Del52_ expressing cells. Ba/F3-MPL cells were treated with Bafilomycin A1 (BafA1), an inhibitor of lysosomal acidification. BafA1 binds to and inhibits the translocation of H^+^ ions through vacuolar H^+^-ATPase (V-ATPase) and results in the inactivation of low pH-dependent lysosomal hydrolases. This reduces endosomal maturation and inhibits lysosomal degradation of proteins ^37, 38^. Inhibition of lysosomal degradation by BafA1 rescued surface MPL levels measured by flow cytometry in both vector control (EV) as well as CRT_Del52_ expressing Ba/F3-MPL cells (**Figures 4A-B**). However, the rescue of surface MPL upon BafA1 treatment was significantly higher in Ba/F3-MPL cells expressing CRT_Del52_ in comparison to EV cells (**Figure 4C**). We further examined a possible additional role of proteasomal degradation in downregulation of surface MPL levels in Ba/F3-MPL cells expressing CRT_Del52_. Ba/F3-MPL cells were treated with proteasomal degradation inhibitors, MG132 and Bortezomib. Both MG132 and bortezomib bind to and inhibit the 20S proteolytic core complex of 26S proteasome. Compared to the untreated condition, both EV and CRT_Del52_ expressing cells exhibited a trend towards an increase in the levels of surface MPL, particularly at the lower concentrations of MG132 (0.5 µM to 10 µM) (**Figure 4D**) and upon bortezomib treatment (**Figure 4E**). However, the effects did not reach significance when compared to untreated cells, and no significant differences were observed in EV *vs*. CRT_Del52_ cells (**Figures 4D- E**).

**Figure 4:**
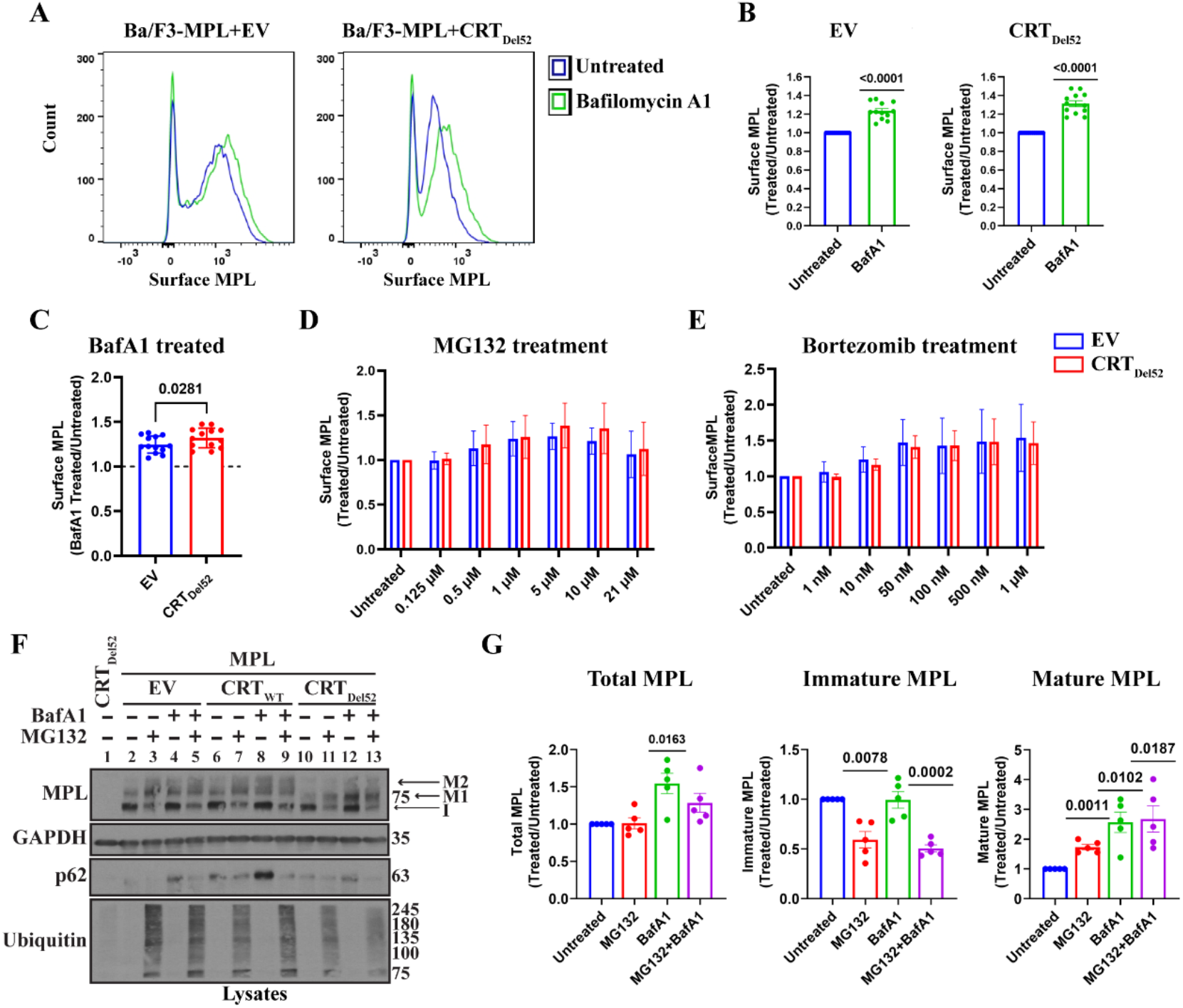
Inhibition of lysosomal acidification rescues cell surface levels of MPL protein more significantly in the presence of CRT_Del52_: (A-C) Murine Ba/F3-MPL control (EV) cells or those expressing CRT_Del52_ were treated with BafilomycinA1 (100 nM) for 4h in media with IL3. Untreated cells are included for comparison. Surface MPL levels were detected by flow cytometry. (A) Representative histograms of surface MPL levels with and without BafilomycinA1 treatment in both the cell lines. (B) Averaged data for surface MPL levels after BafilomycinA1 (BafA1) treatment plotted as a ratio to the levels in the untreated Ba/F3 cells. One sample t-tests are used for determining the statistical significance. (C) Comparisons of BafA1-mediated rescue of surface MPL in control (EV) vs. CRT_Del52_ cells. The p value is calculated using a paired t-test analysis. Panels B and C include data from 12 experiments from 4 independent transductions of BaF3-MPL cells. **(D and E)** Ba/F3-MPL control (EV) and Ba/F3-MPL CRT_Del52_ cells were treated with inhibitors of proteasomal degradation, MG132 (D) or Bortezomib (E) at the indicated concentrations for 4h at 37 °C in media with IL3. Surface MPL levels measured by flow cytometry are plotted as a ratio of MFI values in treated relative to untreated cells (n=3). **(F)** Representative blots (n=5) for MPL in the lysates of Ba/F3-MPL (EV), Ba/F3-MPL-CRT_WT_ or Ba/F3-MPL-CRT_Del52_ cells as indicated treated with 21µM MG132, 100 nM BafA1 or both. The immature form of MPL protein (I) and the two distinct mature forms, M1 (partially mature MPL expressed in CRT_Del52_ cells) and M2 (detected in CRT_WT_ expressing cells), are indicated within the MPL blot. Ubiquitin and p62 were probed as markers to show successful inhibition of proteasomal and lysosomal degradation, respectively. **(G)** Bar graphs show quantifications of total, immature and mature MPL protein in Ba/F3-MPL CRT_Del52_ cells treated with MG132 and/or BafA1 normalized to the values on untreated cells based on immunoblots. Image J was used for quantification of blots. One sample t-tests are used for determining the statistical significance. Graphs were plotted using GraphPad Prism. EV, Empty Vector; IL3, Interleukin 3

The rescue of surface MPL upon BafA1 treatment was also reflected by an increase in the mature MPL fraction in the lysates of Ba/F3-MPL CRT_Del52_ as well as control cells, based on immunoblots (**Figure 4F**, lanes 2 compared to 4, 6 compared to 8 and 10 compared to 12 and **Figure 4G**). On the other hand, 21 µM MG132 treatment induced a rescue of mature MPL along with a parallel reduction in the immature MPL levels (band indicated as ‘**I**’ in **Figure 4F**, lanes 2 compared to 3, 6 compared to 7 and 10 compared to 11; **Figure 4G** and see also **Figure S1**) in all the cells. These effects are observed at the higher concentrations of MG132 or bortezomib (**Figure S1**) and may correspond to forced MPL exit from the ER and glycan maturation under conditions where ER-associated degradation is blocked via proteasome inhibition. Nonetheless, no significant difference in total MPL or cell surface MPL is observed following proteasome inhibition (**Figures 4D, 4E and 4G**).

Overall, the analyses of Figure 4 suggest a dominant role of lysosomal degradation as a regulator of cell surface MPL levels in cells expressing CRT_Del52_. The analyses also show that cells co-expressing CRT_Del52_ and MPL have a lower molecular weight glycoform of mature MPL (**Figure 4F**, lanes 10-13, band labeled M1) compared with EV or CRT_WT_ expressing cells (**Figure 4F**, lanes 2-9, band labeled M2), consistent with previous findings that interaction of CRT_Del52_ with MPL prevents the maturation of N117-linked glycan in MPL ^16^.

### Inhibiting lysosomal degradation rescues CRT_Del52_ levels

To determine the effects of protein degradation pathways on cellular CRT_Del52_ levels in Ba/F3 cells, we treated the cells with MG132,100 nM BafA1 or both. Under all conditions, flow cytometric analyses using anti-CRT(C_mut_) antibody ^17^ indicated low to undetectable surface expression of CRT_Del52_ in the absence of MPL. On the other hand, cell surface CRT_Del52_ is better detectable in cells expressing MPL (**Figures 5A and 5B**), but neither MG132 nor BafA1 rescued surface CRT_Del52_ expression. However, as observed for MPL (Figure 4), BafA1 treatment markedly rescues cellular CRT_Del52_ levels in Ba/F3-MPL cells when compared to untreated cells (CRT_Del52_ blot in **Figure 5C**, lanes 9 *vs.* 11). Some rescue of CRT_Del52_ levels was also observed in the Ba/F3 cells in the absence of MPL co- expression (CRT_Del52_ blot in **Figure 5C**, lanes 5 *vs.* 7), however, the rescue of CRT_Del52_ levels was significant in the cells co-expressing MPL (**Figure 5D**). On the other hand, treatment with MG132 (21 µM) promoted CRT_Del52_ degradation independently of MPL co- expression (CRT_Del52_ blot in **Figure 5C**, lanes 5 *vs.* 6 and lanes 9 *vs.*10). A small but nonsignificant rescue of CRT_Del52_ levels was observed in Ba/F3-MPL cells upon combining the MG132 (21 µM) and BafA1 (100 nM) treatments, compared with MG132 alone (**Figure 5D**).

**Figure 5:**
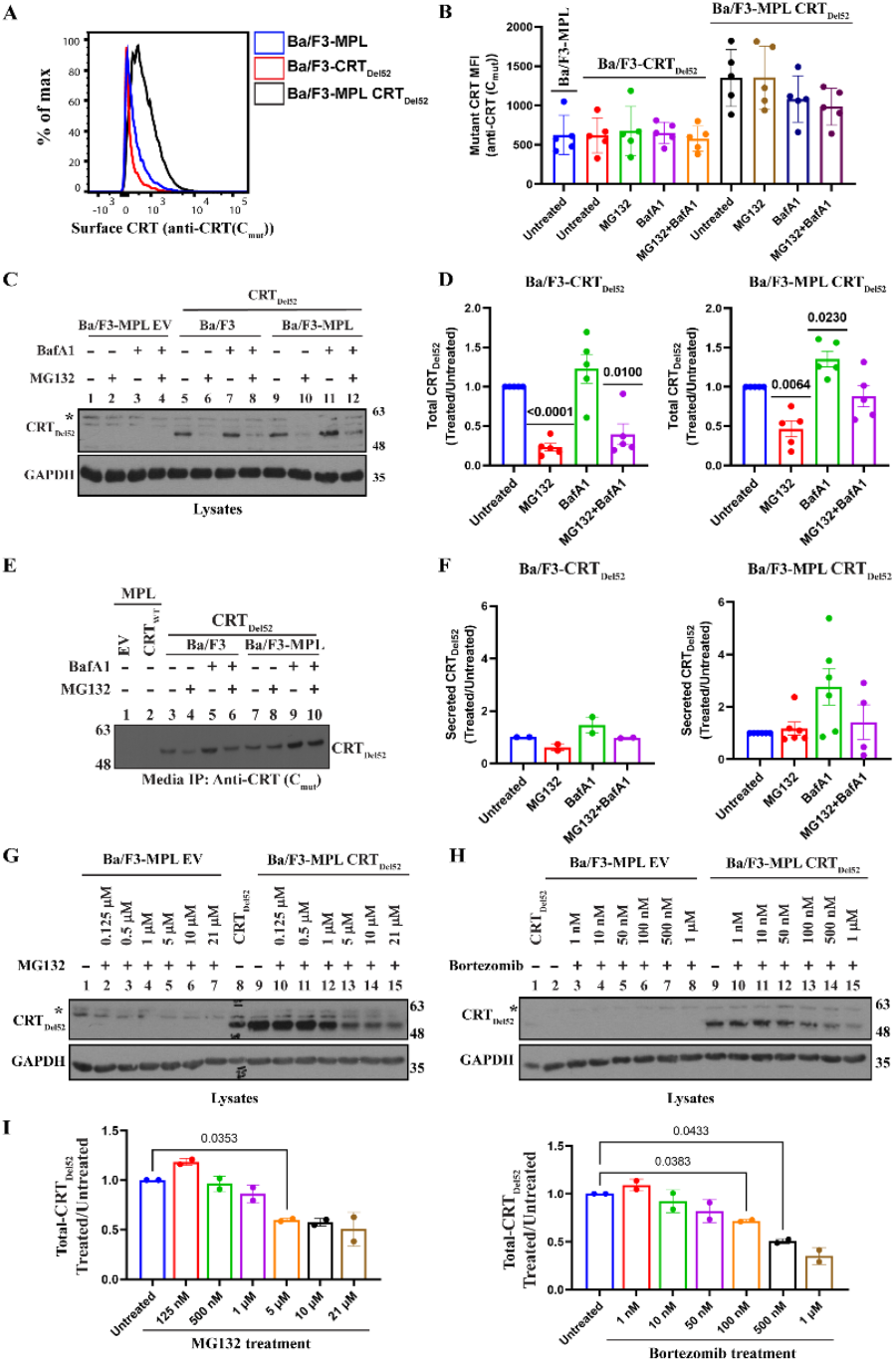
Inhibition of lysosomal degradation rescues CRT_Del52_ levels whereas the proteasomal pathway inhibition promotes CRT_Del52_ degradation: (A and B) Representative histograms (A) and bar graph (B) show mean fluorescence intensity (MFI) of surface CRT detected by flow cytometry using mutant CRT specific antibody (anti-CRT (C_mut_)) on Ba/F3-MPL, Ba/F3-CRT_Del52_ and Ba/F3-MPL CRT_Del52_ cells that were either untreated or treated with 21 µM MG132 and/or 100 nM Bafilomycin A1 (BafA1). **(C and D)** Ba/F3-MPL, Ba/F3-CRT_Del52_ or Ba/F3-MPL CRT_Del52_ cells as indicated were treated with 21 µM MG132 and/or 100 nM Bafilomycin A1 (BafA1) for 4 h at 37°C in media with IL3. The representative blots (n=4) show levels of total CRT_Del52_ (C). Graphs show band intensities of CRT_Del52_ quantified from the blots in treated Ba/F3-CRT_Del52_ or Ba/F3-MPL CRT_Del52_ cells normalized to untreated cells (D). **(E)** Representative blot of secreted CRT_Del52_ immunoprecipitated (IP) using anti-CRT(C_mut_) antibody from cell culture media of Ba/F3-CRT_Del52_ cells (n=2) and Ba/F3-MPL CRT_Del52_ cells (n=6) that were either untreated or treated with 21 µM MG132 and/or 100 nM BafA1 for 4h at 37°C in media with IL3. **(F)** Quantification of secreted CRT_Del52_ band intensities from immunoblots normalized to the values for untreated cells. **(G-I)** Representative immunoblots (n=3) show CRT_Del52_ levels (G and H) in the lysates of control (EV) or CRT_Del52_ expressing Ba/F3-MPL cells treated for 4 h with different concentrations of MG132 (G) or Bortezomib (H). Bar graphs (I) show quantifications of CRT_Del52_ band intensities from blots of Ba/F3-MPL CRT_Del52_ cells (n=2) treated with different concentrations of MG132 or Bortezomib normalized to CRT_Del52_ band intensities in untreated cells. CRT_Del52_ in C, E, G and H were probed using anti-CRT(C_mut_) antibody. GAPDH blots in panels C, G and H show equal loading of lysates in different lanes. Image J was used for quantification of blots. Graphs in panels B, D, F, and I were plotted using GraphPad Prism. Statistical significance indicated by p values was determined using a one-sample t-test in panels D and F and repeated measures one-way ANOVA in panel I.

To compare the effects of the degradation inhibitors upon CRT_Del52_ secretion, we immunoprecipitated CRT_Del52_ from the culture supernatant of Ba/F3 cells following treatments with MG132 and/or BafA1 using the anti-CRT(C_mut_) antibody ^17^. Ba/F3-MPL (EV) and Ba/F3-MPL cells co-expressing wild type CRT (CRT_WT_) were included as controls (**Figure 5E**, lanes 1 and 2). As shown by the representative blots, the absence of bands following IP from control cells (**Figure 5E**, lanes 1 and 2) demonstrated the specificity of IP of CRT_Del52_ from Ba/F3 cells expressing CRT_Del52_. BafA1 treatment resulted in a trend towards rescue of secreted CRT_Del52_ in culture supernatant of Ba/F3 cells irrespective of MPL co-expression (**Figure 5E**, lanes 5 and 9 and **Figure 5F**). MG132 treatment resulted in loss of secreted CRT_Del52_, particularly in the absence of MPL (**Figures 5E and 5F**), and the combination of BafA1 treatment with MG132 showed a small but nonsignificant rescue of secreted CRT_Del52_ in culture supernatant of Ba/F3-MPL cells compared to untreated cells (**Figure 5F**). Together, these results indicate that the inhibition of lysosomal degradation rescues cellular CRT_Del52_ and to some extent the secreted CRT_Del52_.

We treated Ba/F3-MPL CRT_Del52_ cells with different concentrations of proteasomal degradation inhibitors MG132 and bortezomib as discussed for MPL (Figure 4). There is a small non-significant increase in the amounts of CRT_Del52_ in Ba/F3-MPL cells treated with low MG132 and bortezomib concentrations (**Figures 5G and 5H**, lanes 9 compared to 10 and **Figure 5I**). However, CRT_Del52_ levels notably decreased upon treatment with higher concentrations of MG132 (1 µM to 21 µM) (**Figures 5G and 5I**) and bortezomib (50 nM to 1 µM) (**Figures 5H and 5I**) with the levels of CRT_Del52_ being the lowest upon treatment with the highest concentrations of MG132 (21 µM) and bortezomib (1 µM) tested. Thus, inhibition of proteasomal degradation at the higher concentrations of drugs promotes CRT_Del52_ degradation in Ba/F3-MPL cells.

### mTOR inhibition reduces cytokine-independent proliferation mediated by CRT_Del52_

MPN-linked CRT mutants support cytokine-independent proliferation of Ba/F3 cells that co-express MPL ^11^. Figures 4 and 5 show that CRT_Del52_ and MPL are degraded via the lysosomal pathway more significantly in the presence of one another, and inhibition of lysosomal degradation rescues both proteins. Based on these results, we examined whether direct activation of lysosomal degradation using FDA approved clinical drugs could promote CRT_Del52_ and MPL degradation and interfere with the cell proliferative effects of CRT_Del52._ For this, we used Rapamycin and Everolimus, both of which are the inhibitors of mammalian target of rapamycin (mTOR), a protein kinase that inhibits autophagy and promotes cell growth and anabolism in response to several growth factors, nutrients, and other factors ^39^. Ba/F3-MPL EV cells cultured in the presence of IL3 generally show normal growth patterns upon treatment with either 50 nM or 100 nM rapamycin (**Figures 6A and 6B**) or 100 nM everolimus (**Figures 6C and 6D**). A low inhibition of proliferation of Ba/F3-MPL EV cells was measured at early time points upon treatment with rapamycin or everolimus (**Figures 6B and 6D**) which could result from the inhibition of IL3-dependent signaling pathway by mTOR inhibitors ^40^. However, the EV cells recover from the effects of mTOR inhibitors at later time points (**Figures 6B and 6D**). In contrast, Ba/F3-MPL CRT_Del52_ cells show reduced proliferation in the absence of IL3 upon treatment with rapamycin (**Figures 6E and 6F**) and everolimus (**Figures 6G and 6H**). A significant reduction of cytokine-independent proliferation was observed after treatment of Ba/F3-MPL CRT_Del52_ cells with rapamycin starting at Day 2 (**Figure 6F**). A significant reduction of cytokine-independent proliferation of Ba/F3-MPL CRT_Del52_ cells treated with 100 nM everolimus was observed starting even at Day 1 (**Figure 6H**).

**Figure 6:**
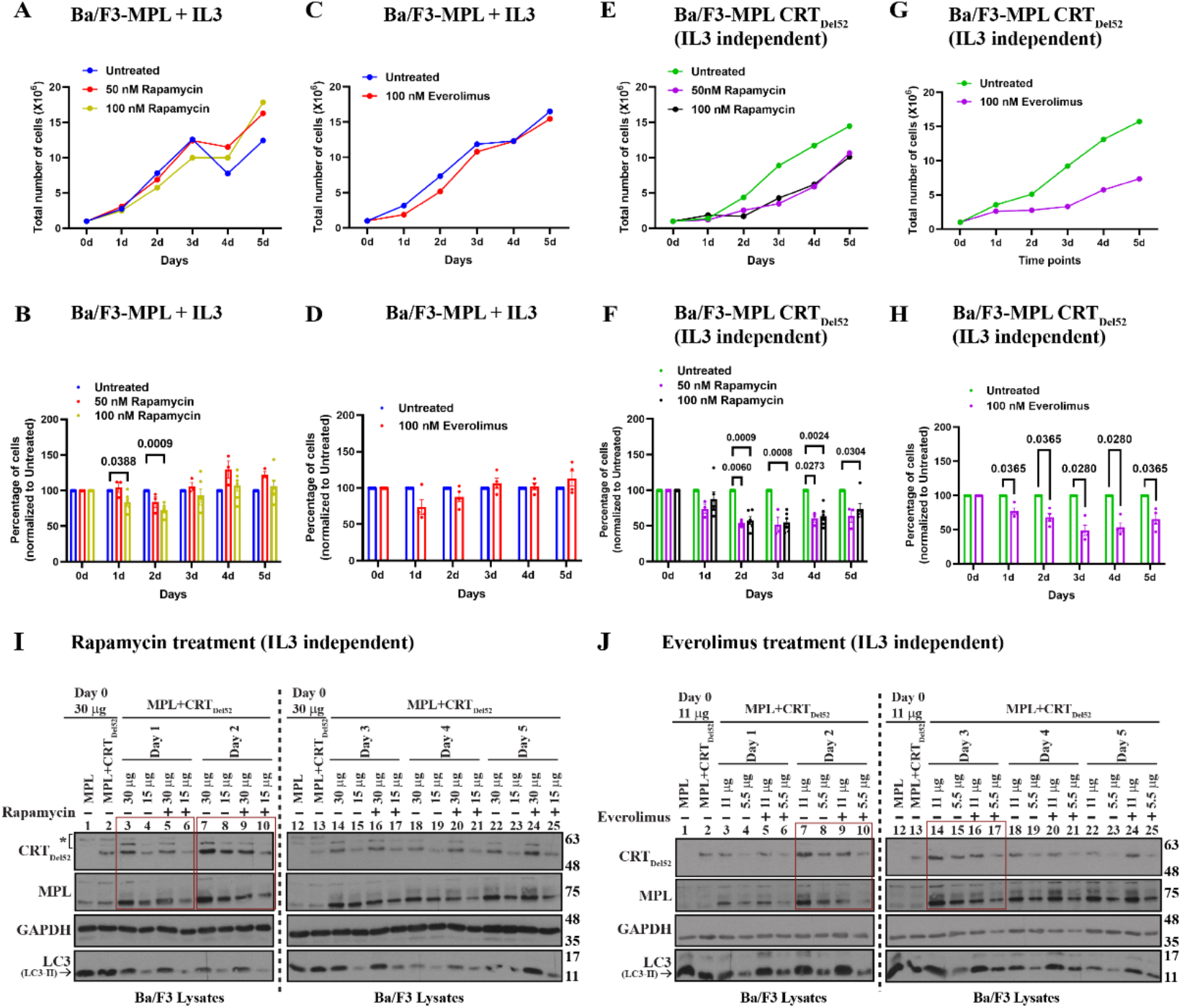
mTOR inhibitors reduce cytokine-independent proliferation mediated by CRT_Del52_. (A, C, E and G) Representative proliferation curves of Ba/F3-MPL (A and C) or Ba/F3-MPL-CRT_Del52_ (E and G) cells that are treated with or without 50 nM (n=3) or 100 nM Rapamycin (n=6) (A and E), with or without 100 nM Everolimus (n=4) (C and G), either in the presence of IL-3 (A and C) or in its absence (IL3-independent; E and G) as indicated. **(B, D, F and H)** Averaged relative proliferation corresponding to the conditions of A, C E and G. Cell counts of the untreated cells were set at 100% for all the time points. Two-way ANOVA analyses (B and F) and multiple paired t-test (D and H) were used to determine statistical significance (indicated by p-values) on different days. **(I and J)** Representative blots of lysates of Ba/F3-MPL CRT_Del52_ cells treated with 100 nM rapamycin (I) or 100 nM everolimus (J) that were harvested at different time points during proliferation assays and compared to the untreated cells. Ba/F3-MPL lysates were loaded as control in the specified lanes. Two serial dilutions of each lysate were loaded in consecutive lanes as indicated. CRT_Del52_ and MPL blots were probed with anti-CRT(C_mut_) and anti-MPL antibodies. GAPDH blots show relative loading of lysates as indicated. LC3 blots show autophagy activation upon Rapamycin treatment marked by LC3-II bands (arrow).

To examine whether reduced proliferation of Ba/F3-MPL cells treated with rapamycin and everolimus is mediated by enhanced degradation of MPL and/or CRT_Del52_, we measured cellular CRT_Del52_ and MPL levels in the lysates of untreated or treated cells at different time points. Immunoblots confirmed that at least at earlier time points (lanes highlighted by boxes), both CRT_Del52_ and MPL levels are lower in cells treated with rapamycin (**Figure 6I**, lanes 3 to 6 and lanes 7 to 10 corresponding to lysates from days 1 and 2, respectively, after the start of drug treatments) and everolimus (**Figure 6J**, lanes 7 to 10 and lanes 14 to 17 corresponding to lysates from days 2 and 3, respectively, after the start of drug treatments) compared to untreated cells. Thus, reduction in cellular CRT_Del52_ and MPL levels triggered by treatment with mTOR-inhibiting (lysosomal degradation-activating) drugs could in part explain the observed reduction in cytokine-independent proliferation of Ba/F3-MPL CRT_Del52_ cells in the presence of rapamycin and everolimus.

### Primary CD34^+^ cells from MPN patients demonstrate lower colony-forming capacity following treatment with everolimus

The results from Figure 6 clearly demonstrate the reduction of cytokine-independent proliferation of Ba/F3-MPL CRT_Del52_ cells upon treatment with mTOR inhibitors, rapamycin and everolimus (Figures 6A-H). To check the susceptibility of primary CD34^+^ cells from CRT-mutated MPN patients to mTOR inhibition, CD34^+^ cells isolated from bone marrow samples of MPN patients (details of patient samples given in Supplemental Table 3) or from healthy donor leukopaks were plated on the semi-solid methylcellulose media for colony forming unit (CFU) assays. In the first set of experiments, the CD34^+^ cells were either left untreated or pre-treated with everolimus for 24 h before plating (**Figure 7A**, pre-treatment). In subsequent experiments, everolimus was incorporated within the plating media (**Figure 7B**, continuous treatment). We used human methylcellulose enriched media (R&D systems; HSC005) that is optimized for growth of myeloid, erythroid, and mixed lineage (colony forming unit-granulocyte, erythrocyte, monocyte, and megakaryocyte (CFU-GEMM)) progenitor cells. In general, the number of CFUs grown from MPN patient CD34^+^ cells were higher than the number of CFUs grown from healthy donor CD34^+^ cells (**Figures 7A and 7B**). Compared to colony counts under untreated conditions, MPN patient CD34^+^ cells exhibited a decreasing trend in the number of CFUs following pre-treatment with 50 nM everolimus (**Figure 7A**, n=6). Under more stringent conditions involving continuous exposure to 50 nM everolimus (**Figure 7B**, n=3), a decrease in the number of CFUs derived from MPN patient CD34^+^ cells was also observed, compared to untreated cells. There was significant heterogeneity in the measured response which included a heterogenous group of available patient samples in different disease states (Supplemental Table 3). Strikingly however, healthy donor CD34^+^ cells exhibited no to little effects of pre-treatment (n=4) or continuous treatment (n=3) with 50 nM everolimus when compared to CFUs derived from untreated healthy CD34^+^ cells (**Figures 7A and 7B**). Thus, despite expected variabilities in primary human samples, the results from figures 6 and 7 clearly indicate that mutant CRT expressing cells or primary CD34^+^ cells from MPN patients are susceptible to the drug-mediated activation of autophagy/lysosomal pathway, unlike healthy donor CD34^+^ cells.

**Figure 7:**
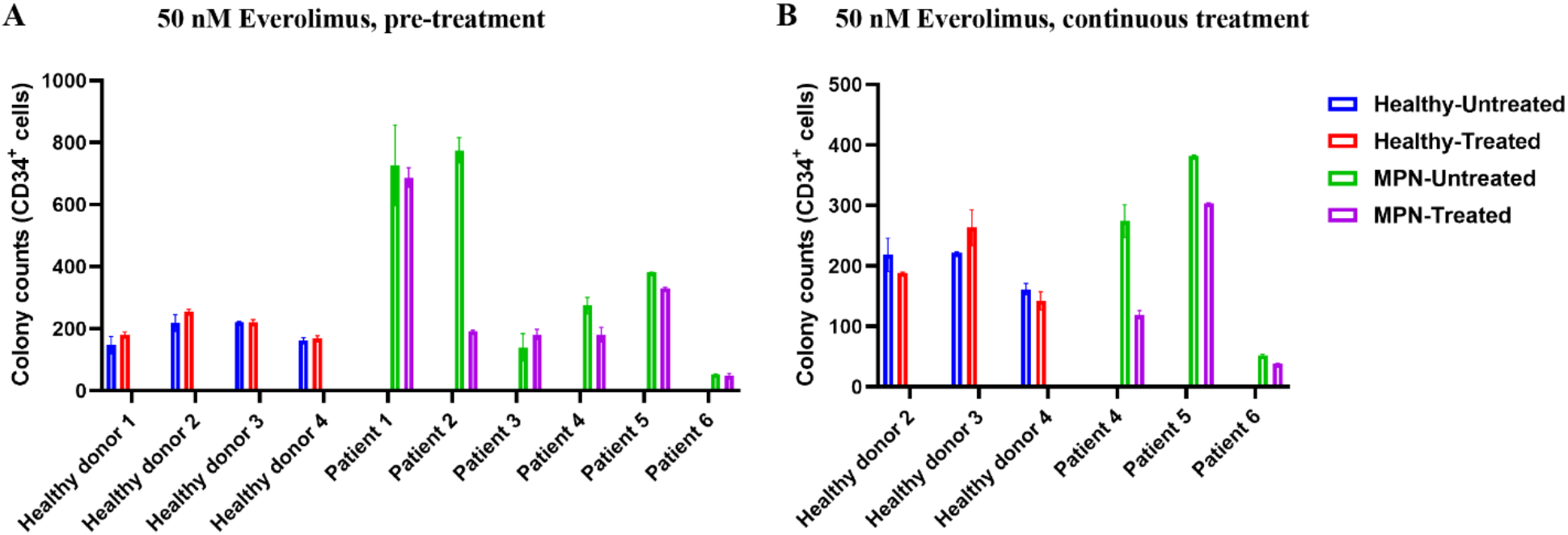
Primary CD34^+^ cells from MPN patients exhibit lower colony-forming capacity following treatment with everolimus. CD34^+^ hematopoietic stem cells (HSCs) isolated from bone marrow samples of MPN patients or mobilized leukopaks from healthy donors were plated on semi-solid methylcellulose containing media. The number of colonies were counted 12-18 days after plating. **(A)** Absolute number of colonies obtained upon culturing MPN patient (six patients) or healthy donor (four donors) CD34^+^ HSCs in semi-solid media that were either untreated or pre-treated with 50 nM everolimus (pre-treatment). **(B)** Absolute number of colonies obtained upon culturing MPN patient (three patients) or healthy donor (three donors) CD34^+^ HSCs in semi-solid media with or without 50 nM everolimus (continuous treatment).

## Discussion

Our results indicate a significant increase in localization of CRT mutants on the surface of both the resting (CD41^+^) and activated platelets (CD62p^+^) isolated from MPN patients when compared to healthy donor platelets (**Figures 1D-G**), consistent with previous findings in other cells ^8–10, 16^. While early studies have reported downregulation of surface MPL on platelets in ET, PV, and MF patients ^41, 42^, whether patients with CRT mutations also have low MPL levels was unknown. Our results indicate that platelets from most of the MPN patients with CRT mutations exhibit low MPL/TPOR levels when compared to healthy donor platelets (**Figure 2**). Both surface MPL and total MPL protein expression are lower in platelets isolated from MPN patient blood samples compared to platelets from healthy donors (**Figures 2B, 2D and 2E**). These findings prompted studies of the role of proteasomal and lysosomal degradation pathways in regulating the dynamics of MPL and mutant CRT proteins in MPNs. Low surface and mature MPL levels measured in Ba/F3 cells expressing both MPL and CRT_Del52_ confirmed that the observed MPL downmodulation is more acute in the cells expressing mutant CRT proteins (**Figure 3**).

Internalization of MPL/TPO complex allows termination of downstream signaling by removing the receptor and other signaling protein complexes from the surface and targeting them towards degradation. MPL internalization occurs via clathrin-dependent endocytosis, requiring residues and motifs within the intracellular domain of MPL ^27, 29^. Mature MPL is suggested to be targeted for degradation by both the lysosomal and proteasomal degradation pathways in response to TPO ^29, 30^. On the other hand, MPL downmodulation in Ba/F3-MPL cells expressing the JAK2V617F mutant was attributed to increased ubiquitination and degradation of the receptor by the proteasomal degradation pathway ^43^. By analogy to these prior findings, we suggest a model wherein low MPL levels on the cell surface in mutant CRT expressing cells result from MPL signaling activated by mutant CRT proteins followed by internalization and lysosomal degradation of MPL-mutant CRT complexes. Notably, lysosomal degradation rather than proteasomal degradation of these complexes is prominently induced in the context of the CRT mutants. In support of this model (**Figure 8**), we find that inhibition of lysosomal acidification markedly rescues cellular CRT_Del52_ levels, particularly in the presence of MPL (**Figures 5C and 5D**). Additionally, lysosomal degradation of cell-surface MPL becomes more prominent in Ba/F3 cells when CRT_Del52_ is co-expressed (**Figure 4C**), and synergistic rescue of both MPL and CRT_Del52_ levels is observed upon inhibition of lysosomal degradation.

**Figure 8:**
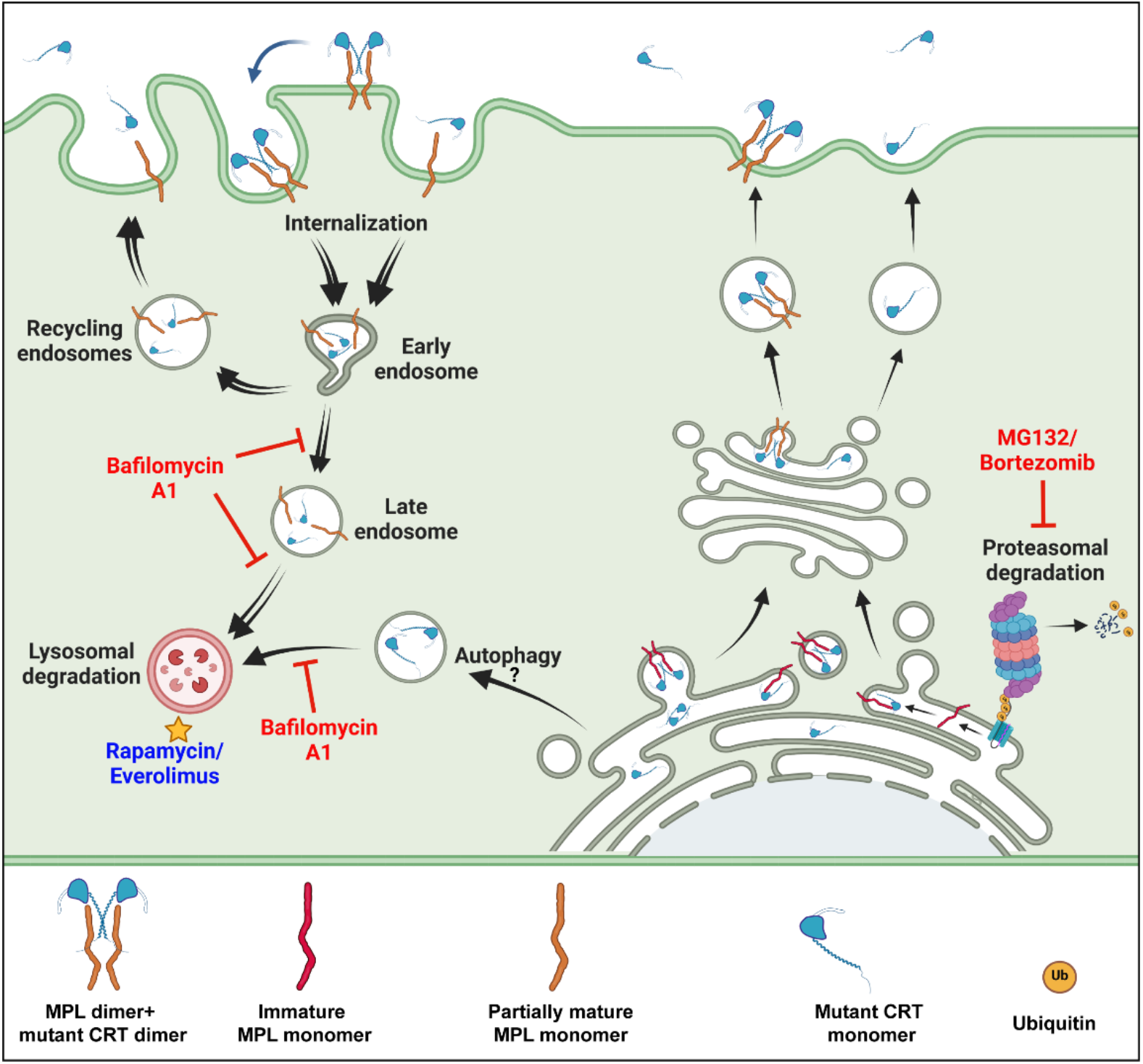
Lysosomal degradation pathways regulate mutant CRT and cell surface MPL levels in mutant CRT-driven MPNs: Mutant CRT is trafficked to cell surface and media as free protein or in complex with MPL, which prevents full glycan maturation on MPL. Treatment with the vacuolar proton-ATPase inhibitor, bafilomycin A1, rescues cellular and secreted mutant CRT protein levels. Surface MPL is also rescued by treatment with bafilomycin, the extent of which is enhanced by the presence of mutant CRT. Bafilomycin-mediated rescue of mutant CRT and MPL is expected to result from inhibition of lysosomal degradation and increased recycling of the endocytosed proteins. mTOR inhibitors, everolimus and rapamycin, induce loss of MPL and mutant CRT proteins, likely via their enhanced lysosomal degradation, and suppress cytokine-independent proliferation triggered by mutant CRT. This pathway could be further exploited for therapeutic interventions in mutant CRT-driven MPNs. In contrast to the effects of bafilomycin, MG132 and bortezomib enhance MPL maturation and mutant CRT degradation, likely via multiple compensatory mechanisms that are induced upon inhibition of proteasomal degradation.

In contrast to the measured accumulation of cellular CRT_Del52_ and cell-surface MPL following treatment with BafA1, the inhibitor of lysosomal degradation, drug-mediated inhibition of proteasomal degradation reduces cellular CRT_Del52_ levels while not increasing secreted mutant calreticulin levels (**Figures 5C-F**). This observation suggests that proteasomal inhibitors enhance degradation of CRT_Del52._ These findings are consistent with the observed decrease in mutant CRT protein levels following inhibition of proteasomal degradation reported in an earlier study ^7^. Reduced CRT_Del52_ levels following inhibition of proteasomal degradation (**Figures 5C, 5D and 5G-I**) is likely caused by enhanced CRT_Del52_ degradation via activation of autophagy/lysosomal pathway ^44^. Recent studies have shown that the proteasomal pathway is upregulated in mutant CRT expressing cells from MPN patients ^45^. Treatment with the proteasomal inhibitor, bortezomib, inhibits the proliferation of mutant CRT expressing HSCs and megakaryocytes ^45, 46^, which is attributed to the combined targeting of the proteasome and an ER stress response prevalent in cells expressing CRT_Del52_. While these studies did not show the effect of proteasomal inhibitors on the expression levels of CRT mutants, it is likely that the proliferation-suppressive effects of the proteasomal inhibitors are, at least in part, driven via their ability to induce CRT_Del52_ degradation.

In contrast to MPL protein levels, expression of MPN-linked mutant CRT proteins has been associated with increased MPL transcription ^7, 47^. In one of these studies, enhanced MPL transcription has been attributed to enhanced binding of ERp57 and transcription factor Fli1 on MPL promoter in CRT_Del52_ expressing cells ^47^. While MPL protein levels were not measured in those studies, the data in **Figure 2** indicate a dominant loss of MPL expression in platelets despite any gain in mRNA expression, further supporting the model that protein degradation contributes to the low measured MPL protein levels. Downregulation of MPL protein levels despite increased MPL mRNA expression was also observed in Ba/F3-MPL cells expressing the JAK2V617F mutant ^43^.

mTOR activity plays a crucial role in regulating lysosomal function by regulating lysosomal biogenesis, lysosomal positioning, and activity of lysosomal proteins ^48^. mTOR inhibition has been correlated to enhanced lysosomal degradation via different mechanisms including enhanced lysosomal biogenesis ^49^ or enhanced lysosomal acidification through assembly of active V-ATPase at lysosomal membranes ^50^. mTOR inhibitors, rapamycin and everolimus, inhibit proliferation and survival of cancer cells ^51, 52^. We further show here that treatments with these drugs, suppress cytokine-independent proliferation of CRT_Del52_ expressing Ba/F3-MPL cells (**Figures 6E-H**). The drugs also induce a decrease in the levels of CRT_Del52_ and MPL readily detectable at early time points after drug treatment (**Figures 6I and 6J**). We also observe a decrease in the number of CFUs recovered from CD34^+^ cells isolated from MPN patients following treatment with everolimus (**Figure 7**). Despite all the variabilities between samples, everolimus treatment had no measured impact on colony growth from CD34^+^ cells of healthy donors (**Figure 7**).

Overall, our findings demonstrate the importance of cellular degradation pathways in regulation of MPL and mutant CRT protein levels, which are the two major factors controlling oncogenesis in mutant CRT-linked MPNs. Everolimus (or RAD001) has been approved as an anti-cancer drug for treatment of breast cancer, renal cell carcinoma and certain types of pancreatic and lung cancers, etc. (www.cancer.gov/about-cancer/treatment/drugs/everolimus). Some studies have indicated mTOR pathway inhibitors as potential drugs for treatment of MPNs ^53, 54^. The combination of an mTOR inhibitor, BEZ235 and JAK2 inhibitor, Ruxolitinib, showed synergistic effects in the treatment of JAK2V617F mice models of MPNs ^55^. Our studies suggest that mTOR inhibitors are promising targets for treatment of MPN patients expressing CRT mutants and highlight their therapeutic potential.

## Materials and methods

### Materials

All common laboratory chemicals were purchased from Sigma Aldrich. The QuikChange site-directed mutagenesis kit was purchased from Agilent (200521). Antibiotic-antimycotic (15240062), Dulbecco’s modified eagle’s medium (DMEM; 11965-092), fetal bovine serum (FBS; 16140071), L-Glutamine (25030081), RPMI 1640 media (11875093), Geneticin (10131035) and Puromycin (A113803) were purchased from Gibco (ThermoFisher Scientific). Lipofectamine LTX (15338100) was procured from ThermoFisher Scientific. Fugene® HD reagent was purchased from Promega (E2311). Apyrase (A6535), Acid-citrate dextrose solution (C3821), MG132 (474787) and Protease Inhibitor Cocktail (P8340) were obtained from Sigma Aldrich. Bafilomycin (11038) was procured from Cayman Chemicals. Drugs Rapamycin (AY22989), Everolimus (S1120) and Bortezomib (S1013) were purchased from Selleck Chemicals. Human Methylcellulose enriched media (HSC005) was purchased from R&D systems. The Endo H (P0703) enzyme was purchased from NEB. Recombinant mouse IL3 (575506) was procured from BioLegend. PE-conjugated anti-MPL (562159) used for flow cytometry was obtained from BD biosciences. EasySep™ human CD34 positive selection kit II was procured from Stem cell technologies (17856). Ficoll^®^ Paque PLUS density gradient media was purchased from Sigma Aldrich (GE17-1440-03).

### Study Approval

Blood samples were collected after obtaining written informed consent from donors and MPN patients following protocols approved by the University of Michigan Institutional Review Board. MPN patient blood samples were collected from the Myeloproliferative diseases repository (study ID: HUM0006778). Healthy donor blood samples were obtained from the University of Michigan Platelet Physiology and Pharmacology Core repository (study ID: HUM00107120) and from participants of another research study (study ID: HUM00071750).

### DNA Constructs

The constructs of wild type CRT and MPN-linked CRT mutants used for expression in mammalian cells have been described earlier ^17^. Untagged human MPL construct (accession number BC153092) was purchased from DNASU (clone ID: HsCD00350714) and subcloned into pMSCV-neomycin retroviral vector at EcoRI and XhoI sites. All CRT and MPL primers used are specified in Supplemental Table 1.

### Cell lines and retroviral transduction

Human embryonic kidney (HEK) 293T cells were cultured in DMEM supplemented with 10% FBS and antibiotic-antimycotic (1X). The mouse pro B cell line, Ba/F3, was maintained in RPMI 1640 supplemented with mouse IL-3 (10 ng/mL), 10% FBS and antibiotic-antimycotic (1X). Wherever mentioned, puromycin and neomycin were added to media for culture of Ba/F3 cells at 3 µg/mL and 1000 µg/mL, respectively.

Ba/F3 cells were transduced with DNA constructs mentioned above for stable CRT or MPL expression. HEK293T cells (∼70% confluent at 1X10^6^ cells per well of a six-well plate) were transfected with 2.75 µg of pMSCV constructs of MPL or CRT along with pCL-Eco (2 µg) and VSV-G (0.25 µg) vectors expressing gag/pol and envelope proteins, respectively, using Fugene® HD reagent. Viral supernatants were collected after 48 hours, filtered using 0.45 µm nylon membrane syringe filters, and used for transduction. Ba/F3 cells were first transduced to express human MPL (Ba/F3-MPL cells) followed by a second round of transduction for expression of either human CRT_WT_ or CRT_Del52_ protein. Empty vector controls (EV) were included. Transduction was performed by spinoculation for 2 hours at 2500 rpm at room temperature (RT) by adding viral supernatant collected from one well of a six-well plate to 0.5x10^6^ Ba/F3 cells in the presence of 8 µg/mL polybrene. Following spinoculation, two-thirds of the media volume was replaced with fresh media containing IL3. Cells were harvested after 24 hours to completely remove viral supernatant-containing media. Antibiotic selection was started 48 hours after spinoculation. Protein expression was checked following antibiotic selection using protocols described in the immunoblotting section.

For drug treatment experiments, cells were incubated for four hours in RPMI media with mouse IL-3 and specific concentrations of MG132, Bortezomib or Bafilomycin (BafA1).

### Custom anti-CRT mutant C-terminus-specific antibody production

As described earlier ^17^, a mutant C-terminus specific anti-CRT(C_mut_) antibody was custom generated by GeneTel Laboratories following immunization of rabbits with a 22-mer peptide (KMSPARPRTSCREACLQGWTEA) corresponding to the unique sequence at carboxy-termini of mutant CRT proteins.

### Isolation of Platelets

Whole blood samples from de-identified patients or healthy donors were received in BD Vacutainer^®^ ACD solution A tubes (catalog number: 364606). Platelet isolation was performed as described previously ^56^ with some modifications. For the preparation of platelet-rich plasma (PRP), blood samples were centrifuged in polypropylene tubes on a swing bucket rotor at slow speed (200 x g) for 15 min at room temperature (RT) using slow acceleration and zero deceleration settings. PRP was transferred to fresh polypropylene tubes and treated with 0.02 U/mL apyrase and 1 mL ACD solution (C3821 Sigma) for every 9 mL of PRP. Platelets were isolated from PRP by centrifugation at 2000 x g for 10 min at RT after adjusting acceleration/deceleration to low. The platelets were resuspended in Tyrode’s buffer containing 10 mM HEPES pH 7.4, 134 mM NaCl, 12 mM NaHCO_3_, 2.9 mM KCl, 0.34 mM Na_2_HPO_4_, 1 mM MgCl_2_ and 5 mM D-glucose and rested for 30 min at 37°C. Platelets were either stained as described in the flow cytometry section or pelleted at 2000 x g followed by lysis for 30 min at 4°C in lysis buffer (composition given in immunoblotting section). The lysates were used for immunoblotting or stored at -80°C.

### Cell lysis, Endo H digestion and immunoblotting

Ba/F3 cells or platelets from MPN patients and healthy donors were lysed in lysis buffer (50 mM Tris pH 7.5, 150 mM NaCl, 1% Triton X-100, 5 mM CaCl_2_ and protease inhibitor cocktail (P8340 from Sigma @ 1:100 dilution)) for 30 min at 4°C. Total protein was quantified in the lysates using BCA reagent kit. For Endo H digestion, lysate volume equivalent to 20-40 µg total protein was first denatured at 100°C for 10 min in 1X glycoprotein denaturation buffer (NEB). Endo H digestion was performed at 37°C for 1h in the presence of 1X GlycoBuffer 3 and 1 µL of Endo H enzyme (NEB). The lysates were boiled and subjected to SDS-PAGE and western transfer. Antibodies used for immunoblotting were: anti-CRT(N) antibody (Cell Signaling Technology (CST) 12238, 1:10,000 dilution) for detection of CRT_WT_; anti-CRT(C_mut_) antibody for detection of CRT mutants; anti-MPL antibody (EMD Millipore 06944, 1:2500 dilution) for detection of MPL; anti- SQSTM1/p62 (CST 5114, 1:10000 dilution) and anti-LC3 (Novus biologicals NB100- 2220, 1:10000 dilution) antibodies for detection of p62 and LC3, respectively; anti-ubiquitin (CST 3936, 1:10000 dilution) and anti-GAPDH (CST 2118, 1:20,000 dilution) antibodies for detection of ubiquitin and GAPDH, respectively. Blots were incubated with Horseradish peroxidase-conjugated secondary antibodies (Jackson Immunoresearch) and developed via chemiluminescence (Pierce™ ECL western blotting substrate, 32106).

Image J was used for the quantification of band intensities on western blots.

### Flow Cytometry

For surface staining, unfixed platelets were incubated with appropriate primary antibody cocktail for 30 min in staining buffer (PBS + 2% FBS) at RT, followed by two washes with staining buffer. Platelets were then incubated in appropriate dilution of secondary antibody in staining buffer for 15 min at RT and washed twice with staining buffer. For total staining, platelets were first fixed with 4% paraformaldehyde (PFA) in PBS for 10 min at RT followed by incubation with appropriate dilution of primary antibodies in staining buffer with 0.2% saponin for 30 min at RT. Wherever applicable, appropriate dilution of Alexa Fluor™ conjugated secondary antibody in staining buffer with 0.2% saponin was added to platelets for 15 min at RT. Staining parameters were recorded on BD LSRFortessa cell analyzer.

CD41 and CD62p proteins on platelets were stained with FITC conjugated anti-CD41 (BioLegend 303704,1:50 dilution) and PE conjugated anti-CD62p (BD Pharmingen 561921, 1:50 dilution) antibodies, respectively. Surface CRT was detected using anti-CRT(N) or anti-CRT(C_mut_) antibody (1:200 dilution) followed by staining with Alexa Fluor™ 647 conjugated secondary antibody (Invitrogen A21245, 1:2000 dilution). Total CRT in healthy donor platelets was stained with anti-CRT(N) primary antibody and Alexa Fluor™ 647 conjugated secondary antibody. Surface or total MPL proteins were stained with anti-human MPL-PE antibody (BD Pharmingen 562159; 1:50 dilution).

Ba/F3 cells expressing human MPL protein that were either cultured normally or treated with drugs were harvested and stained for surface MPL using PE-conjugated anti-human MPL antibody (BD Pharmingen 562159,1:50 dilution) in staining buffer for 30 min at 4°C.

For total MPL staining, Ba/F3 cells were first stained with live/dead™ fixable aqua dead cell stain (Invitrogen L34957, 1:1000 dilution) for 10 min in 1X PBS at RT, followed by fixation in 4% PFA for 10 min at RT. Total MPL was then stained using a PE-conjugated anti-human MPL antibody (BD Pharmingen) in staining buffer with 0.2% saponin for 30 min at 4°C.

FlowJo™ was used for analysis of flow cytometry data.

### Immunoprecipitation (IP) of secreted forms of CRT_Del52_

Approximately 5 million Ba/F3 cells expressing MPL alone or those expressing CRT_WT_ or CRT_Del52_ protein with or without MPL co-expression were incubated in a six-well plate in 4 mL media with IL3 in the presence of either MG132 (10 μg/mL or 21 µM) and/or BafA1 (100 nM) or no drug (untreated) for 4 hours at 37°C. Cells along with culture media were harvested and centrifuged at high speed (∼14000 X g) for 5 min to remove cells and the culture media was saved and filtered to remove any remaining cells. Secreted forms of mutant CRT proteins were immunoprecipitated from filtered media using anti-CRT(C_mut_) antibody (at 2 μg/mL final antibody concentration). The IP was performed overnight at 4°C followed by incubation with protein G beads (at ∼ 20 μl beads/mL of culture supernatant) for 2 hours at 4°C to harvest immune complexes. Beads were washed with lysis buffer (composition given in the immunoblotting section) and boiled in SDS loading dye to elute immune complexes bound to beads. Samples were subjected to immunoblotting as mentioned above.

### Proliferation assays

Ba/F3-MPL EV cells and Ba/F3-MPL CRT_Del52_ cells were seeded in the presence and absence of mouse IL3, respectively, on day 0 at a density of 1 million cells/well in a six-well plate in 3 mL RPMI media per well. Cells were either left untreated or treated with 50 nM or 100 nM Rapamycin or 100 nM Everolimus. Cells were counted after trypan blue staining on each day for the following five days (day 1 to day 5). On days 3, 4 and 5, fresh media with or without drug was supplemented in all wells (same volume of media was added in all the wells) to prevent cell death due to high confluency. Total number of cells in untreated or treated wells counted on each day were plotted to generate the proliferation assay graphs in GraphPad Prism 9.

For detection of CRT_Del52_ levels by immunoblotting, cells were seeded in separate wells for each time point during proliferation assay. At each time point, cells were counted, harvested, washed and cell pellets were stored at -80°C until lysis. All the cell pellets were lysed at the same time after the collection of cells on day 5 time point.

### Colony forming unit (CFU) assays with CD34^+^ cells

For CFU assays with CD34^+^ cells, bone marrow samples from MPN patients or leukopaks from healthy donors were used (Supplemental Table 3). Frozen bone marrow cells from MPN patients or frozen leukopaks from healthy donors were thawed and incubated overnight in RPMI media with DNase 1 (Roche 10104159001, 0.1 mg/mL). Freshly collected bone marrow samples from some MPN patients were used directly for cell isolation. The cell suspension was filtered through 70 µm cell strainer(s) to remove any clumps and subjected to density gradient centrifugation on Ficoll^®^ Paque Plus media. Cells in the buffy coat were harvested, washed to remove Ficoll, counted and used for isolation of CD34^+^ cells using EasySep™ human CD34 positive selection kit II (Stem cell technologies) following manufacturer’s protocol. For untreated condition or continuous treatment with everolimus, ∼20,000 CD34^+^ cells were incubated in duplicates for 24 hours in 2 mL RPMI+10% FBS media without any drugs. For pre-treatment condition, 50 nM everolimus was added to the cell suspension during the 24-hour incubation prior to plating. After 24 hours, cells from each well were harvested and resuspended in 100 µL of cell resuspension solution provided with human methylcellulose enriched media (R&D systems). Each cell suspension was added to 2.5 mL methylcellulose enriched media, vortexed and rested to allow bubbles to escape. For continuous treatment, 50 nM everolimus was added into the methylcellulose media at this step. Cells were plated in methylcellulose media in a twelve-well plate and incubated for 12-18 days at 37°C until colonies were of optimum size for counting. Colony counts were determined manually or using Image J for some experiments and plotted using GraphPad Prism.

### Statistical analysis

All statistical analysis was performed in GraphPad Prism 9.

## Author Contributions

AK and AV designed and performed experiments, analyzed data, wrote the original draft and edited the manuscript. MK collected patient blood and bone marrow samples, purified platelets and edited the manuscript. MT is the director of MPN repository at the University of Michigan. MR designed and supervised the study, obtained funding, analyzed data, and wrote, and edited the manuscript.

## Acknowledgements

We are grateful to all the donors and patients who volunteered to donate blood and/or bone marrow samples for this work. We thank Amanda Prieur from the University of Michigan Platelet Physiology and Pharmacology Core for collecting healthy donor blood samples. We thank Polk Avery from Talpaz lab for coordinating and collecting MPN patient blood and bone marrow samples. This work was funded by National Institute of Health grants (R01 AI123957) to MR and the University of Michigan Fast Forward Protein Folding Diseases Initiative.

## Supplemental data

**Figure S1:**
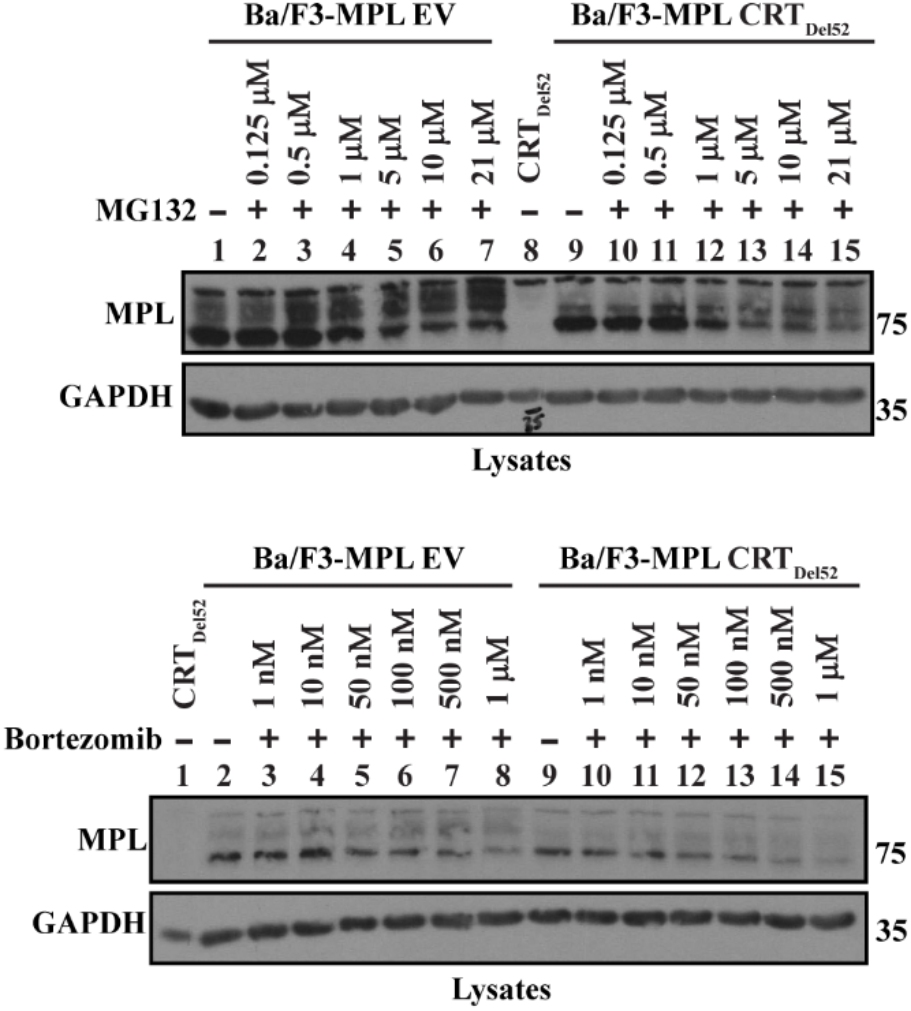
Representative blots (n=2) of Ba/F3-MPL and Ba/F3-MPL-CRT_Del52_ cell lysates after treatment with indicated concentrations of MG132 or Bortezomib for 4 h at 37 °C in media with IL3. GAPDH blots show equal loading of lysates.

**Supplemental Table 1:**
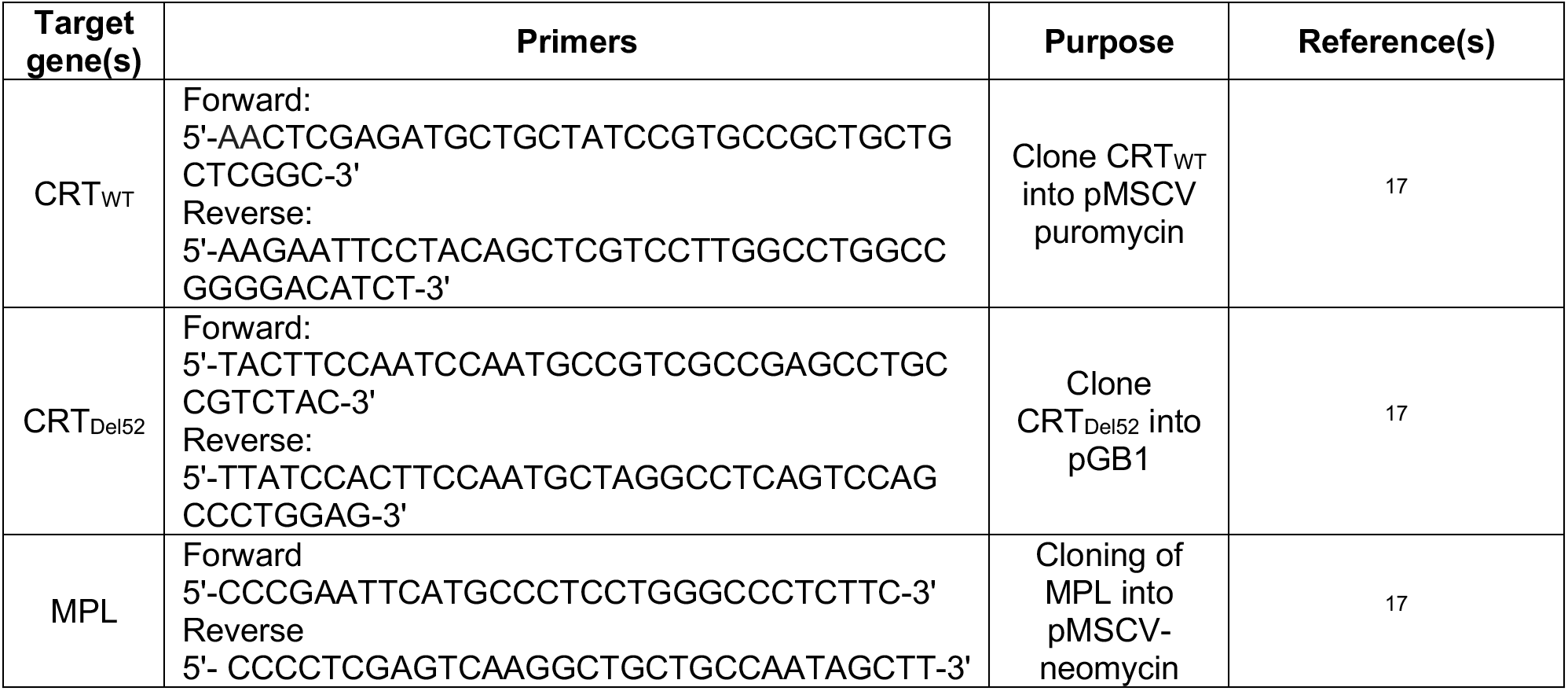
List of primers used in this study

**Supplemental Table 2:**
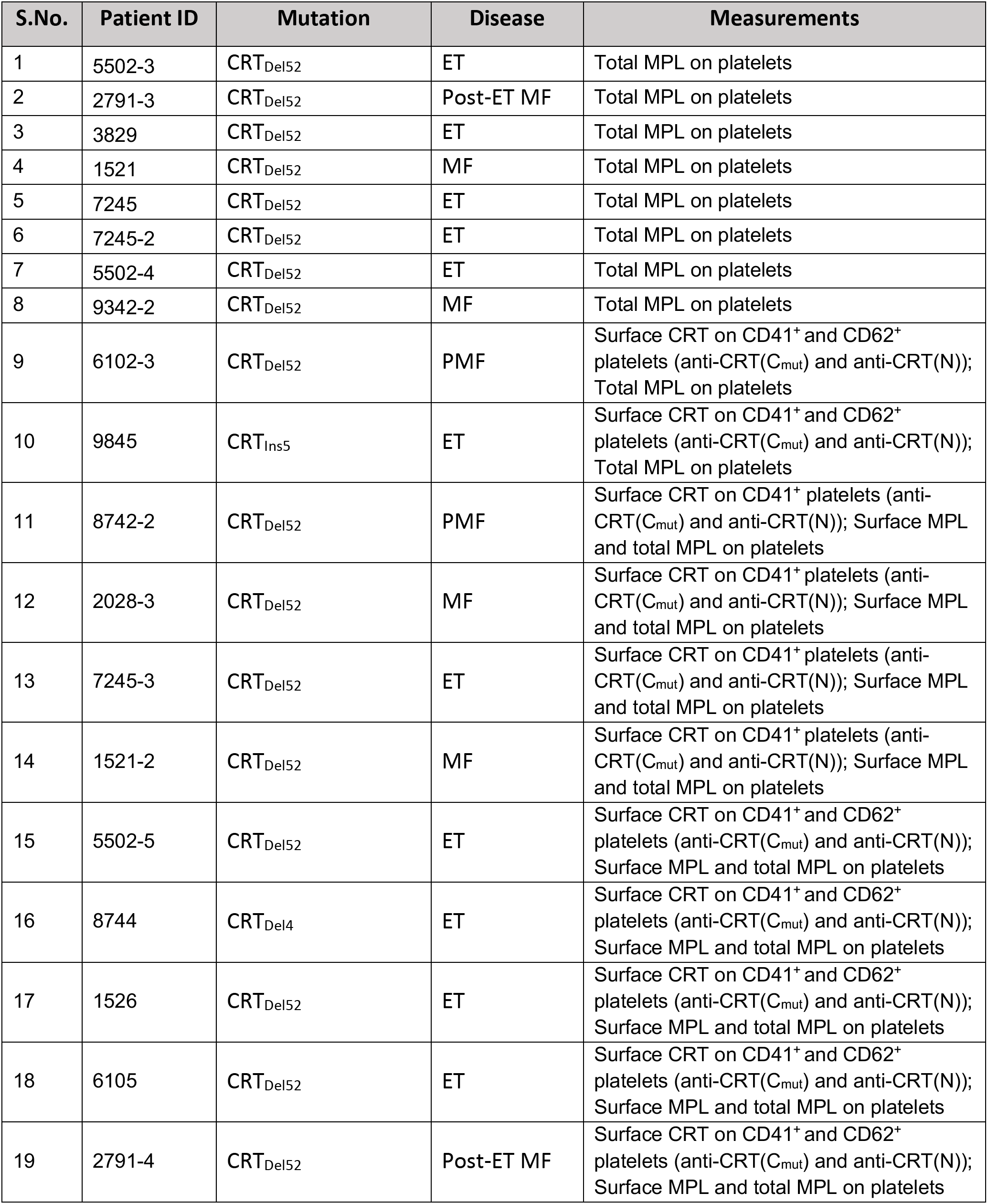
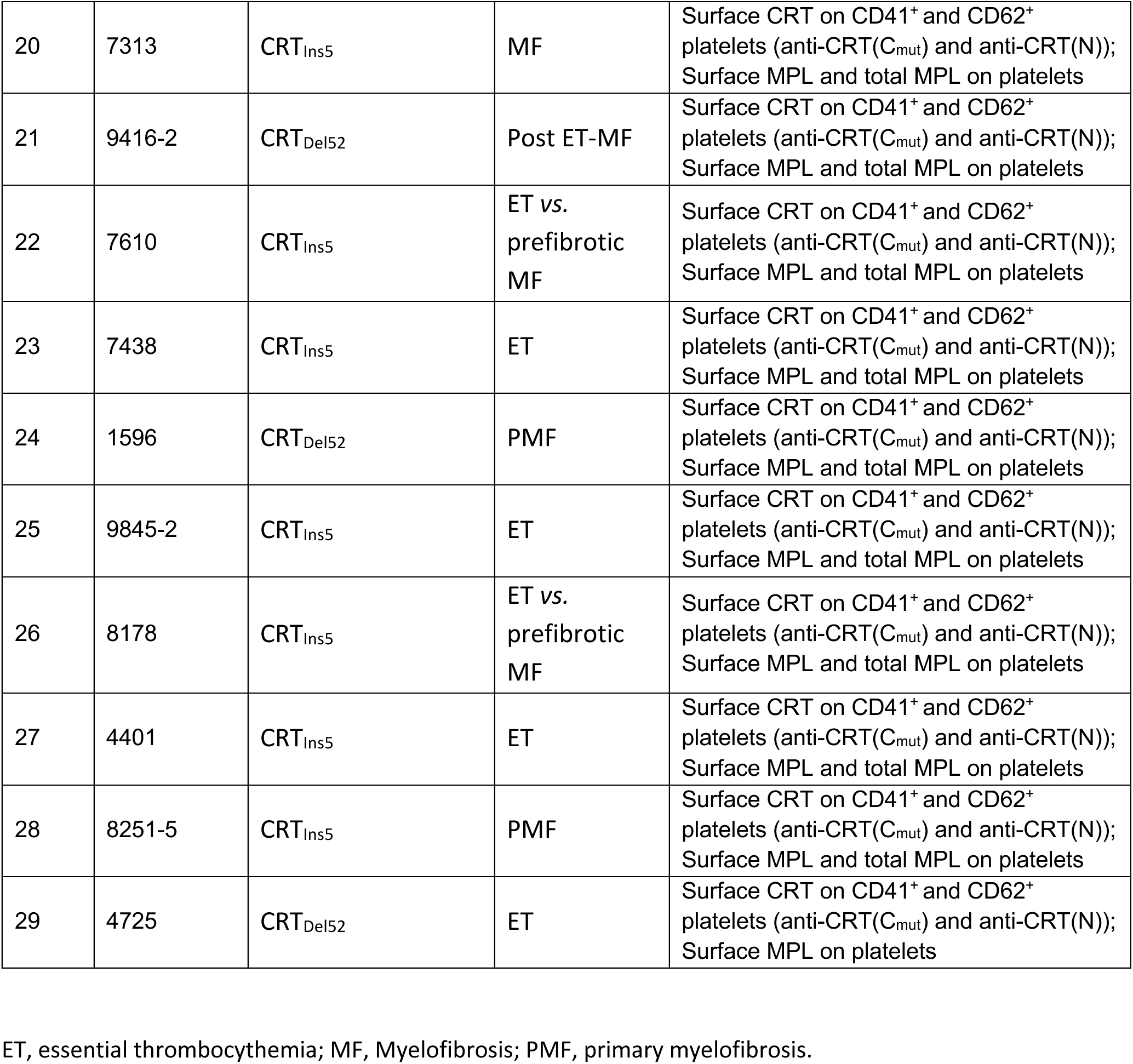
List of MPN patient samples used for surface CRT and/or surface and total MPL measurements in platelets

| S.No. | Patient ID | Mutation | Disease | Measurements |
| --- | --- | --- | --- | --- |
| 1 | 5502-3 | CRT<sub>Del52</sub> | ET | Total MPL on platelets |
| 2 | 2791-3 | CRT<sub>Del52</sub> | Post-ET MF | Total MPL on platelets |
| 3 | 3829 | CRT<sub>Del52</sub> | ET | Total MPL on platelets |
| 4 | 1521 | CRT<sub>Del52</sub> | MF | Total MPL on platelets |
| 5 | 7245 | CRT<sub>Del52</sub> | ET | Total MPL on platelets |
| 6 | 7245-2 | CRT<sub>Del52</sub> | ET | Total MPL on platelets |
| 7 | 5502-4 | CRT<sub>Del52</sub> | ET | Total MPL on platelets |
| 8 | 9342-2 | CRT<sub>Del52</sub> | MF | Total MPL on platelets |
| 9 | 6102-3 | CRT<sub>Del52</sub> | PMF | Surface CRT on CD41<sup>+</sup> and CD62<sup>+</sup> platelets (anti-CRT(C<sub>mut</sub>) and anti-CRT(N)); Total MPL on platelets |
| 10 | 9845 | CRT<sub>Ins5</sub> | ET | Surface CRT on CD41<sup>+</sup> and CD62<sup>+</sup> platelets (anti-CRT(C<sub>mut</sub>) and anti-CRT(N)); Total MPL on platelets |
| 11 | 8742-2 | CRT<sub>Del52</sub> | PMF | Surface CRT on CD41<sup>+</sup> platelets (anti-CRT(C<sub>mut</sub>) and anti-CRT(N)); Surface MPL and total MPL on platelets |
| 12 | 2028-3 | CRT<sub>Del52</sub> | MF | Surface CRT on CD41<sup>+</sup> platelets (anti-CRT(C<sub>mut</sub>) and anti-CRT(N)); Surface MPL and total MPL on platelets |
| 13 | 7245-3 | CRT<sub>Del52</sub> | ET | Surface CRT on CD41<sup>+</sup> platelets (anti-CRT(C<sub>mut</sub>) and anti-CRT(N)); Surface MPL and total MPL on platelets |
| 14 | 1521-2 | CRT<sub>Del52</sub> | MF | Surface CRT on CD41<sup>+</sup> platelets (anti-CRT(C<sub>mut</sub>) and anti-CRT(N)); Surface MPL and total MPL on platelets |
| 15 | 5502-5 | CRT<sub>Del52</sub> | ET | Surface CRT on CD41<sup>+</sup> and CD62<sup>+</sup> platelets (anti-CRT(C<sub>mut</sub>) and anti-CRT(N)); Surface MPL and total MPL on platelets |
| 16 | 8744 | CRT<sub>Del4</sub> | ET | Surface CRT on CD41<sup>+</sup> and CD62<sup>+</sup> platelets (anti-CRT(C<sub>mut</sub>) and anti-CRT(N)); Surface MPL and total MPL on platelets |
| 17 | 1526 | CRT<sub>Del52</sub> | ET | Surface CRT on CD41<sup>+</sup> and CD62<sup>+</sup> platelets (anti-CRT(C<sub>mut</sub>) and anti-CRT(N)); Surface MPL and total MPL on platelets |
| 18 | 6105 | CRT<sub>Del52</sub> | ET | Surface CRT on CD41<sup>+</sup> and CD62<sup>+</sup> platelets (anti-CRT(C<sub>mut</sub>) and anti-CRT(N)); Surface MPL and total MPL on platelets |
| 19 | 2791-4 | CRT<sub>Del52</sub> | Post-ET MF | Surface CRT on CD41<sup>+</sup> and CD62<sup>+</sup> platelets (anti-CRT(C<sub>mut</sub>) and anti-CRT(N)); Surface MPL and total MPL on platelets |
| 20 | 7313 | CRT<sub>Ins5</sub> | MF | Surface CRT on CD41<sup>+</sup> and CD62<sup>+</sup> platelets (anti-CRT(C<sub>mut</sub>) and anti-CRT(N));<br>Surface MPL and total MPL on platelets |
| 21 | 9416-2 | CRT<sub>Del52</sub> | Post ET-MF | Surface CRT on CD41<sup>+</sup> and CD62<sup>+</sup> platelets (anti-CRT(C<sub>mut</sub>) and anti-CRT(N));<br>Surface MPL and total MPL on platelets |
| 22 | 7610 | CRT<sub>Ins5</sub> | ET vs.<br>prefibrotic<br>MF | Surface CRT on CD41<sup>+</sup> and CD62<sup>+</sup> platelets (anti-CRT(C<sub>mut</sub>) and anti-CRT(N));<br>Surface MPL and total MPL on platelets |
| 23 | 7438 | CRT<sub>Ins5</sub> | ET | Surface CRT on CD41<sup>+</sup> and CD62<sup>+</sup> platelets (anti-CRT(C<sub>mut</sub>) and anti-CRT(N));<br>Surface MPL and total MPL on platelets |
| 24 | 1596 | CRT<sub>Del52</sub> | PMF | Surface CRT on CD41<sup>+</sup> and CD62<sup>+</sup> platelets (anti-CRT(C<sub>mut</sub>) and anti-CRT(N));<br>Surface MPL and total MPL on platelets |
| 25 | 9845-2 | CRT<sub>Ins5</sub> | ET | Surface CRT on CD41<sup>+</sup> and CD62<sup>+</sup> platelets (anti-CRT(C<sub>mut</sub>) and anti-CRT(N));<br>Surface MPL and total MPL on platelets |
| 26 | 8178 | CRT<sub>Ins5</sub> | ET vs.<br>prefibrotic<br>MF | Surface CRT on CD41<sup>+</sup> and CD62<sup>+</sup> platelets (anti-CRT(C<sub>mut</sub>) and anti-CRT(N));<br>Surface MPL and total MPL on platelets |
| 27 | 4401 | CRT<sub>Ins5</sub> | ET | Surface CRT on CD41<sup>+</sup> and CD62<sup>+</sup> platelets (anti-CRT(C<sub>mut</sub>) and anti-CRT(N));<br>Surface MPL and total MPL on platelets |
| 28 | 8251-5 | CRT<sub>Ins5</sub> | PMF | Surface CRT on CD41<sup>+</sup> and CD62<sup>+</sup> platelets (anti-CRT(C<sub>mut</sub>) and anti-CRT(N));<br>Surface MPL and total MPL on platelets |
| 29 | 4725 | CRT<sub>Del52</sub> | ET | Surface CRT on CD41<sup>+</sup> and CD62<sup>+</sup> platelets (anti-CRT(C<sub>mut</sub>) and anti-CRT(N));<br>Surface MPL on platelets |

ET, essential thrombocythemia; MF, Myelofibrosis; PMF, primary myelofibrosis.

**Supplemental Table 3.** List of MPN patient bone marrow samples used for CD34^+^ CFU assays

| S.No. | Patient ID | Mutation | Disease | Treatments for CFU assays |
| --- | --- | --- | --- | --- |
| 1 | **0542** | CRT<sub>Del52</sub> | MF | Pre-treatment with 50 nM Everolimus |
| 2 | **7245** | CRT<sub>Del52</sub> | ET | Pre-treatment with 50 nM Everolimus |
| 3 | **3124** | CRT<sub>Del52</sub> | ET | Pre-treatment with 50 nM Everolimus |
| 4 | **2791** | CRT<sub>Del52</sub> | Post-ET MF | Pre-treatment with 50 nM Everolimus;<br>continued treatment with 50 nM Everolimus |
| 5 | **1086** | CRT<sub>Del52</sub> | PMF | Pre-treatment with 50 nM Everolimus;<br>continued treatment with 50 nM Everolimus |
| 6 | **0266** | CRT<sub>Del52</sub> | Post-ET MF | Pre-treatment with 50 nM Everolimus;<br>continued treatment with 50 nM Everolimus |

ET, essential thrombocythemia; MF, Myelofibrosis; PMF, primary myelofibrosis.

## References

[1] Campbell PJ, Green AR: The Myeloproliferative Disorders. New England Journal of Medicine 2006, 355:2452–66.

[2] Spivak JL: Myeloproliferative Neoplasms. New England Journal of Medicine 2017, 376:2168–81.

[3] Klampfl T, Gisslinger H, Harutyunyan AS, Nivarthi H, Rumi E, Milosevic JD, Them NCC, Berg T, Gisslinger B, Pietra D, Chen D, Vladimer GI, Bagienski K, Milanesi C, Carola Casetti I, Sant E, Ferretti V, Elena C, Schischlik F, Cleary C, Six M, Schalling M, Schönegger A, Bock C, Malcovati L, Pascutto C, Superti-Furga G, Cazzola M, Kralovics R: Somatic Mutations of Calreticulin in Myeloproliferative Neoplasms. N Engl J Med 2013, 369:2379–90.

[4] Nangalia J, Massie CE, Baxter EJ, Nice FL, Gundem G, Wedge DC, Avezov E, Li J, Kollmann K, Kent DG, Aziz A, Godfrey AL, Hinton J, Martincorena I, Van Loo P, Jones AV, Guglielmelli P, Tarpey P, Harding HP, Fitzpatrick JD, Goudie CT, Ortmann CA, Loughran SJ, Raine K, Jones DR, Butler AP, Teague JW, O’Meara S, McLaren S, Bianchi M, Silber Y, Dimitropoulou D, Bloxham D, Mudie L, Maddison M, Robinson B, Keohane C, Maclean C, Hill K, Orchard K, Tauro S, Du MQ, Greaves M, Bowen D, Huntly BJP, Harrison CN, Cross NCP, Ron D, Vannucchi AM, Papaemmanuil E, Campbell PJ, Green AR: Somatic CALR mutations in myeloproliferative neoplasms with nonmutated JAK2. New England Journal of Medicine 2013, 369:2391–405.

[5] Masubuchi N, Araki M, Yang Y, Hayashi E, Imai M, Edahiro Y, Hironaka Y, Mizukami Y, Kihara Y, Takei H, Nudejima M, Koike M, Ohsaka A: Mutant calreticulin interacts with MPL in the secretion pathway for activation on the cell surface. Leukemia 2020, 34:499–509.

[6] Arshad N, Cresswell P: Tumor-associated calreticulin variants functionally compromise the peptide loading complex and impair its recruitment of MHC-I. Journal of Biological Chemistry 2018, 293:9555–69.

[7] Han L, Schubert C, Köhler J, Schemionek M, Isfort S, Brümmendorf TH, Koschmieder S, Chatain N: Calreticulin-mutant proteins induce megakaryocytic signaling to transform hematopoietic cells and undergo accelerated degradation and Golgi-mediated secretion. Journal of Hematology and Oncology 2016, 9:1–14.

[8] Liu P, Zhao L, Loos F, Marty C, Xie W, Martins I, Lachkar S, Qu B, Waeckel-Énée E, Plo I, Vainchenker W, Perez F, Rodriguez D, López-Otin C, van Endert P, Zitvogel L, Kepp O, Kroemer G: Immunosuppression by Mutated Calreticulin Released from Malignant Cells. Molecular Cell 2020, 77:748–60.e9.

[9] Pecquet C, Papadopoulos N, Balligand T, Chachoua I, Tisserand A, Vertenoeil G, Nédélec A, Vertommen D, Roy A, Marty C, Nivarthi H, Defour J-P, El-Khoury M, Hug E, Majoros A, Xu E, Zagrijtschuk O, Fertig TE, Marta DS, Gisslinger H, Gisslinger B, Schalling M, Casetti I, Rumi E, Pietra D, Cavalloni C, Arcaini L, Cazzola M, Komatsu N, Kihara Y, Sunami Y, Edahiro Y, Araki M, Lesyk R, Buxhofer-Ausch V, Heibl S, Pasquier F, Havelange V, Plo I, Vainchenker W, Kralovics R, Constantinescu SN: Secreted Mutant Calreticulins As Rogue Cytokines in Myeloproliferative Neoplasms. Blood 2023, 141:917–29.

[10] Araki M, Yang Y, Masubuchi N, Hironaka Y, Takei H, Morishita S, Mizukami Y, Kan S, Shirane S, Edahiro Y, Sunami Y, Ohsaka A, Komatsu N: Activation of the thrombopoietin receptor by mutant calreticulin in CALR-mutant myeloproliferative neoplasms. Blood 2016, 127:1307–16.

[11] Chachoua I, Pecquet C, El-Khoury M, Nivarthi H, Albu R-I, Marty C, Gryshkova V, Defour J-P, Vertenoeil G, Ngo A, Koay A, Raslova H, Courtoy PJ, Choong ML, Plo I, Vainchenker W, Kralovics R, Constantinescu SN: Thrombopoietin receptor activation by myeloproliferative neoplasm associated calreticulin mutants. Blood 2016, 127:1325–35.

[12] Elf S, Abdelfattah NS, Chen E, Perales-Patón J, Rosen EA, Ko A, Peisker F, Florescu N, Giannini S, Wolach O, Morgan EA, Tothova Z, Losman J-A, Schneider RK, Al-Shahrour F, Mullally A: Mutant Calreticulin Requires Both Its Mutant C-terminus and the Thrombopoietin Receptor for Oncogenic Transformation. Cancer discovery 2016, 6:368–81.

[13] Marty C, Pecquet C, Nivarthi H, El-Khoury M, Chachoua I, Tulliez M, Villeval J-L, Raslova H, Kralovics R, Constantinescu SN, Plo I, Vainchenker W: Calreticulin mutants in mice induce an MPL-dependent thrombocytosis with frequent progression to myelofibrosis. Blood 2016, 127:1317–24.

[14] Elf S, Abdelfattah NS, Baral AJ, Beeson D, Rivera JF, Ko A, Florescu N, Birrane G, Chen E, Mullally A: Defining the requirements for the pathogenic interaction between mutant calreticulin and MPL in MPN. Blood 2018, 131:782–6.

[15] Papadopoulos N, Nedelec A, Derenne A, Sulea TA, Pecquet C, Chachoua I, Vertenoeil G, Tilmant T, Petrescu AJ, Mazzucchelli G, Iorga BI, Vertommen D, Constantinescu SN: Oncogenic CALR mutant C-terminus mediates dual binding to the thrombopoietin receptor triggering complex dimerization and activation. Nat Commun 2023, 14:1881.

[16] Pecquet C, Chachoua I, Roy A, Balligand T, Vertenoeil G, Leroy E, Albu R-I, Defour J-P, Nivarthi H, Hug E, Xu E, Ould-Amer Y, Mouton C, Colau D, Vertommen D, Shwe MM, Marty C, Plo I, Vainchenker W, Kralovics R, Constantinescu SN: Calreticulin mutants as oncogenic rogue chaperones for TpoR and traffic-defective pathogenic TpoR mutants. Blood 2019, 133:2669–81.

[17] Venkatesan A, Geng J, Kandarpa M, Wijeyesakere SJ, Bhide A, Talpaz M, Pogozheva ID: Mechanism of mutant calreticulin-mediated activation of the thrombopoietin receptor in cancers. Journal of Cell Biology 2021, 220:e202009179-e.

[18] Desikan H, Kaur A, Pogozheva ID, Raghavan M: Effects of calreticulin mutations on cell transformation and immunity. J Cell Mol Med 2023, 27:1032–44.

[19] Debili N, Wendling F, Cosman D, Titeux M, Florindo C, Dusanter-Fourt I, Schooley K, Methia N, Charon M, Nador R, Bettaieb A, Vainchenker W: The Mpl Receptor Is Expressed in the Megakaryocytic Lineage From Late Progenitors to Platelets. Blood 1995, 85:391–401.

[20] Qian H, Buza-Vidas N, Hyland CD, Jensen CT, Antonchuk J, Månsson R, Thoren LA, Ekblom M, Alexander WS, Jacobsen SEW: Critical Role of Thrombopoietin in Maintaining Adult Quiescent Hematopoietic Stem Cells. Cell Stem Cell 2007, 1:671–84.

[21] Wendling F, Maraskovsky E, Debili N, Florindo C, Teepe M, Titeux M, Methia N, Breton-Gorius J, Cosman D, Vainchenker W: c-Mpl ligand is a humoral regulator of megakaryocytopoiesis. Nature 1994, 369:571–4.

[22] Balligand T, Achouri Y, Pecquet C, Gaudray G, Colau D, Hug E, Rahmani Y, Stroobant V, Plo I, Vainchenker W, Kralovics R, Van den Eynde BJ, Defour J-P, Constantinescu SN: Knock-in of murine Calr del52 induces essential thrombocythemia with slow-rising dominance in mice and reveals key role of Calr exon 9 in cardiac development. Leukemia 2020, 34:510–21.

[23] Shide K, Kameda T, Yamaji T, Sekine M, Inada N, Kamiunten A, Akizuki K, Nakamura K, Hidaka T, Kubuki Y, Shimoda H, Kitanaka A, Honda A, Sawaguchi A, Abe H, Miike T, Iwakiri H, Tahara Y, Sueta M, Hasuike S, Yamamoto S, Nagata K, Shimoda K: Calreticulin mutant mice develop essential thrombocythemia that is ameliorated by the JAK inhibitor ruxolitinib. Leukemia 2017, 31:1136–44.

[24] Benlabiod C, Cacemiro MDC, Nedelec A, Edmond V, Muller D, Rameau P, Touchard L, Gonin P, Constantinescu SN, Raslova H, Villeval JL, Vainchenker W, Plo I, Marty C: Calreticulin del52 and ins5 knock-in mice recapitulate different myeloproliferative phenotypes observed in patients with MPN. Nat Commun 2020, 11:4886.

[25] Varghese LN, Defour J-P, Pecquet C, Constantinescu SN: The thrombopoietin receptor: structural basis of traffic and activation by ligand, mutations, agonists, and mutated calreticulin. Frontiers in Endocrinology 2017, 8:59.

[26] Qian S, Fu F, Li W, Chen Q, de Sauvage FJ: Primary Role of the Liver in Thrombopoietin Production Shown by Tissue-Specific Knockout. Blood 1998, 92:2189–91.

[27] Dahlen DD, Broudy VC, Drachman JG: Internalization of the thrombopoietin receptor is regulated by 2 cytoplasmic motifs. Blood 2003, 102:102–8.

[28] Fielder PJ, Gurney AL, Stefanich E, Marian M, Moore MW, Carver-Moore K, de Sauvage FJ: Regulation of Thrombopoietin Levels by c-mpl–Mediated Binding to Platelets. Blood 1996, 87:2154–61.

[29] Hitchcock IS, Chen MM, King JR, Kaushansky K: YRRL motifs in the cytoplasmic domain of the thrombopoietin receptor regulate receptor internalization and degradation. Blood 2008, 112:2222–31.

[30] Saur SJ, Sangkhae V, Geddis AE, Kaushansky K, Hitchcock IS: Ubiquitination and degradation of the thrombopoietin receptor c-Mpl. Blood 2010, 115:1254–63.

[31] Phillips DR, Charo IF, Parise LV, Fitzgerald LA: The Platelet Membrane Glycoprotein IIb-IIIa Complex. Blood 1988, 71:831–43.

[32] Leytin V, Mody M, Semple JW, Garvey B, Freedman J: Flow cytometric parameters for characterizing platelet activation by measuring P-selectin (CD62) expression: theoretical consideration and evaluation in thrombin-treated platelet populations. Biochem Biophys Res Commun 2000, 269:85–90.

[33] McEver RP: GMP-140: a receptor for neutrophils and monocytes on activated platelets and endothelium. Journal of cellular biochemistry 1991, 45:156–61.

[34] Raghavan M, Wijeyesakere SJ, Peters LR, Del Cid N: Calreticulin in the immune system: ins and outs. Trends in Immunology 2013, 34:13–21.

[35] Abbott C, Huang G, Ellison AR, Chen C, Arora T, Szilvassy SJ, Wei P: Mouse Monoclonal Antibodies Against Human c-Mpl and Characterization for Flow Cytometry Applications. Hybridoma 2010, 29:103–13.

[36] Royer Y, Staerk J, Costuleanu M, Courtoy PJ, Constantinescu SN: Janus Kinases Affect Thrombopoietin Receptor Cell Surface Localization and Stability*. Journal of Biological Chemistry 2005, 280:27251–61.

[37] Wang R, Wang J, Hassan A, Lee C-H, Xie X-S, Li X: Molecular basis of V-ATPase inhibition by bafilomycin A1. Nature Communications 2021, 12:1782.

[38] Yamamoto A, Tagawa Y, Yoshimori T, Moriyama Y, Masaki R, Tashiro Y: Bafilomycin A1 prevents maturation of autophagic vacuoles by inhibiting fusion between autophagosomes and lysosomes in rat hepatoma cell line, H-4-II-E cells. Cell Structure and Function 1998, 23:33–42.

[39] Kim YC, Guan K-L: mTOR: a pharmacologic target for autophagy regulation. The Journal of Clinical Investigation 2015, 125:25–32.

[40] Cruz R, Hedden L, Boyer D, Kharas MG, Fruman DA, Lee-Fruman KK: S6 kinase 2 potentiates interleukin-3-driven cell proliferation. Journal of Leukocyte Biology 2005, 78:1378–85.

[41] Horikawa Y, Matsumura I, Hashimoto K, Shiraga M, Kosugi S, Tadokoro S, Kato T, Miyazaki H, Tomiyama Y, Kurata Y, Matsuzawa Y, Kanakura Y: Markedly reduced expression of platelet c-mpl receptor in essential thrombocythemia. Blood 1997, 90:4031–8.

[42] Moliterno AR, Hankins WD, Spivak JL: Impaired expression of the thrombopoietin receptor by platelets from patients with polycythemia vera. N Engl J Med 1998, 338:572–80.

[43] Pecquet C, Diaconu CC, Staerk J, Girardot M, Marty C, Royer Y, Defour JP, Dusa A, Besancenot R, Giraudier S, Villeval JL, Knoops L, Courtoy PJ, Vainchenker W, Constantinescu SN: Thrombopoietin receptor down-modulation by JAK2 V617F: restoration of receptor levels by inhibitors of pathologic JAK2 signaling and of proteasomes. Blood 2012, 119:4625–35.

[44] Li C, Wang X, Li X, Qiu K, Jiao F, Liu Y, Kong Q, Liu Y, Wu Y: Proteasome Inhibition Activates Autophagy-Lysosome Pathway Associated With TFEB Dephosphorylation and Nuclear Translocation. Front Cell Dev Biol 2019, 7:170.

[45] Jutzi JS, Marneth AE, Jimenez-Santos MJ, Hem J, Guerra-Moreno A, Rolles B, Bhatt S, Myers SA, Carr SA, Hong Y, Pozdnyakova O, van Galen P, Al-Shahrour F, Nam AS, Mullally A: CALR-mutated cells are vulnerable to combined inhibition of the proteasome and the endoplasmic reticulum stress response. Leukemia 2023, 37:359–69.

[46] Foßelteder J, Pabst G, Sconocchia T, Schlacher A, Auinger L, Kashofer K, Beham-Schmid C, Trajanoski S, Waskow C, Scholl W, Sill H, Zebisch A, Wolfler A, Thomas D, Reinisch A: Human gene-engineered calreticulin mutant stem cells recapitulate MPN hallmarks and identify targetable vulnerabilities. Leukemia 2023, 37:843–53.

[47] Pronier E, Cifani P, Merlinsky TR, Berman KB, Somasundara AVH, Rampal RK, LaCava J, Wei KE, Pastore F, Maag JL, Park J, Koche R, Kentsis A, Levine RL: Targeting the CALR interactome in myeloproliferative neoplasms. JCI Insight 2018, 3:e122703.

[48] Puertollano R: mTOR and lysosome regulation. F1000Prime Rep 2014, 6:52.

[49] Settembre C, Zoncu R, Medina DL, Vetrini F, Erdin S, Erdin S, Huynh T, Ferron M, Karsenty G, Vellard MC, Facchinetti V, Sabatini DM, Ballabio A: A lysosome-to-nucleus signalling mechanism senses and regulates the lysosome via mTOR and TFEB. EMBO J 2012, 31:1095–108.

[50] Ratto E, Chowdhury SR, Siefert NS, Schneider M, Wittmann M, Helm D, Palm W: Direct control of lysosomal catabolic activity by mTORC1 through regulation of V-ATPase assembly. Nat Commun 2022, 13:4848.

[51] Guertin DA, Sabatini DM: An expanding role for mTOR in cancer. Trends Mol Med 2005, 11:353–61.

[52] Sabatini DM: mTOR and cancer: insights into a complex relationship. Nat Rev Cancer 2006, 6:729–34.

[53] Guglielmelli P, Barosi G, Rambaldi A, Marchioli R, Masciulli A, Tozzi L, Biamonte F, Bartalucci N, Gattoni E, Lupo ML, Finazzi G, Pancrazzi A, Antonioli E, Susini MC, Pieri L, Malevolti E, Usala E, Occhini U, Grossi A, Caglio S, Paratore S, Bosi A, Barbui T, Vannucchi AM, investigators AI-GIMM: Safety and efficacy of everolimus, a mTOR inhibitor, as single agent in a phase 1/2 study in patients with myelofibrosis. Blood 2011, 118:2069–76.

[54] Vannucchi AM, Harrison CN: Emerging treatments for classical myeloproliferative neoplasms. Blood 2017, 129:693–703.

[55] Bartalucci N, Tozzi L, Bogani C, Martinelli S, Rotunno G, Villeval JL, Vannucchi AM: Co-targeting the PI3K/mTOR and JAK2 signalling pathways produces synergistic activity against myeloproliferative neoplasms. J Cell Mol Med 2013, 17:1385–96.

[56] Greening DW, Sparrow RL, Simpson RJ: Preparation of platelet concentrates. Methods Mol Biol 2011, 728:267–78.

